# The Intra-Host Evolutionary Landscape And Pathoadaptation Of Persistent *Staphylococcus aureus* In Chronic Rhinosinusitis

**DOI:** 10.1101/2023.03.28.534496

**Authors:** Ghais Houtak, George Bouras, Roshan Nepal, Gohar Shaghayegh, Clare Cooksley, Alkis James Psaltis, Peter-John Wormald, Sarah Vreugde

**Affiliations:** Adelaide Medical School, Faculty of Health and Medical Sciences, The University of Adelaide, Adelaide, Australia; The Department of Surgery - Otolaryngology Head and Neck Surgery, University of Adelaide and the Basil Hetzel Institute for Translational Health Research, Central Adelaide Local Health Network, South Australia, Australia

## Abstract

Chronic rhinosinusitis (CRS) is a common chronic sinonasal mucosal inflammation associated with *Staphylococcus aureus* biofilm and relapsing infections. This study aimed to determine rates of *S. aureus* persistence and pathoadaptation in CRS patients by investigating the genomic relatedness and antibiotic resistance/tolerance in longitudinally collected *S. aureus* clinical isolates.

A total of 68 *S. aureus* isolates were sourced from 34 CRS patients at least six months apart. Isolates were grown into 48-hour biofilms and tested for tolerance to antibiotics. A hybrid sequencing strategy was used to obtain high-quality reference-grade assemblies of all isolates. Single nucleotide variants (SNV) divergence in the core genome and sequence type clustering were used to analyse the relatedness of the isolate pairs. Single nucleotide and structural genome variations, plasmid similarity, and plasmid copy numbers between pairs were examined.

Our analysis revealed that 41% (14/34 pairs) of *S. aureus* isolates were persisters, while 59% (20/34 pairs) were non-persisters. Persister isolates showed episode-specific mutational changes over time with a bias towards events in genes involved in adhesion to the host and mobile genetic elements such as plasmids, prophages, and insertion sequences. A significant increase in the copy number of conserved plasmids of persister strains (p<0.05) was seen, indicating a role of the “mobilome” in promoting persistence. This was accompanied by a significant increase in biofilm tolerance against all tested antibiotics (p<0.001), which was linked to a significant increase in biofilm biomass (p<0.05) over time, indicating a biofilm central pathoadaptive process in persisters. In conclusion, our study provides important insights into the mutational changes underlying *S. aureus* persistence in CRS patients highlighting pathoadaptive mechanisms in *S. aureus* persisters culminating in increased biofilm biomass linked to an increase in plasmid copy number over time.

## Introduction

Chronic rhinosinusitis (CRS) is characterised by ongoing inflammation of the paranasal sinuses and nasal mucosal lining, which causes symptoms such as nasal congestion, diminished sense of smell, facial pain, and breathing difficulties (Fokkens et al., 2020). Around 10% of people worldwide suffer from CRS, making it a common condition (Hastan et al., 2011).

CRS is clinically subdivided based on its phenotype into two subcategories, CRS with nasal polyps (CRSwNP) and CRS without nasal polyps (CRSsNP) (Hopkins, 2019). Although the pathogenesis of CRS remains unknown, it is known to be a heterogeneous multi-factorial chronic inflammatory disease that frequently co-occurs with conditions such as ciliary dysfunction, aspirin-exacerbated respiratory disease (AERD), and asthma (Fokkens et al., 2020).

It is thought that microbes impact the pathophysiology of CRS. One of the bacteria most abundantly found in the sinuses of CRS patients is *Staphylococcus aureus*, which is frequently associated with exacerbations of the condition (Okifo, Ray, & Gudis, 2022; Vickery, Ramakrishnan, & Suh, 2019).

Several mechanisms of involvement of *S. aureus* in the pathophysiology of CRS have been proposed, including *S. aureus* biofilms as a modulator of chronic mucosal inflammation and relapsing infections (Hoggard et al., 2017; Vickery et al., 2019). Moreover, *S. aureus* mucosal biofilms are associated with poor post-surgical outcomes (Psaltis, Weitzel, Ha, & Wormald, 2008; Singhal, Foreman, Jervis-Bardy, & Wormald, 2011).

Despite the lack of high-level evidence for the effectiveness of antibiotics in treating CRS and its exacerbations, they are commonly prescribed to CRS patients (Barshak & Durand, 2017; Fokkens et al., 2020). Moreover, antibiotics are often ineffective at eliminating the biofilm nidus resulting in a relapsing course of infectious exacerbations (C. W. Hall & Mah, 2017).

Previously we have shown with pulsed-field gel electrophoresis that subjects suffering from CRS are colonised with identical pulsotype *S. aureus* strains months apart in 79% of cases despite multiple courses of systemic antibiotics (Drilling et al., 2014). This suggests that the bacteria can persist in the sinuses despite antibiotic treatment. However, what is less clear is the pathogenic adaptation and phenotypic changes that occur during chronic infection of difficult-to-treat CRS patients.

This study aimed to evaluate the intra-host relatedness of longitudinal *S. aureus* clinical isolates (CI) collected from the nasal cavities of subjects suffering from CRS and characterise the adaption that enables persistence in the host using hybrid long and short read assembled reference-level genomes. Furthermore, intra- and inter-host variability in *S. aureus* phenotype regarding antimicrobial resistance and biofilm tolerance to antibiotics was evaluated to identify phenotypic pathoadaptation of persistent strains.

## Materials and Methods

### Ethics

This project was approved by the Central Adelaide Local Health Network Human Research Ethics Committee under the following reference number: HREC/15/TQEH/132.

### Clinical isolate retrieval

*S. aureus* CIs were retrieved from a bacterial biobank comprised of samples stored in 25% glycerol stock at −80 °C, obtained from swabs taken from the sinonasal cavity of subjects. The swabs were collected from ear-nose-throat inpatient clinic follow-ups and during sinonasal surgery. To be included in this study, longitudinal CI pairs had to be isolated from swabs obtained from patients who fulfilled the EPOS 2020 criteria for difficult-to-treat CRS (Fokkens et al., 2020). The diagnostic criteria and retrieval of asthma, aspirin sensitivity and CRS subtype are elaborated in supplementary text ST1. Only clinical isolate pairs with a time difference of over five months between collections were included in the study. When a subject had more than two clinical isolates available at different timepoints, the isolated pair with the largest time difference was selected. We termed the first recovered isolate group T0, whereas the isolates recovered at later timepoints were termed T1. For all experiments, the clinical isolates were grown overnight on nutrient agar plates (Thermo Fisher Scientific, CM0003, Waltham, USA) from glycerol stock at 37°C unless otherwise specified.

### Antibiotic exposure

The antibiotic exposure of subjects was assessed based on the antibiotic scripts in their medical records. All antibiotic treatments prescribed to the subjects between their first and second sample collection were extracted. The total antibiotic exposure was calculated as the cumulative number of days prescribed for the treatments (Schechner, Temkin, Harbarth, Carmeli, & Schwaber, 2013).

### Genomic DNA extraction and sequencing

For all clinical isolates, hybrid long and short sequencing was performed. The genomic DNA of the *S. aureus* clinical isolates was extracted using the DNeasy Blood & Tissue Kit (Qiagen, 69504, Hilden, Germany) following the manufacturer’s guidelines. The extracted DNA was sequenced using the Oxford nanopore technology (ONT) on the MinION Mk1C (Oxford Nanopore Technologies, Oxford, UK) for long-read sequencing. The Rapid Barcoding Kit (Oxford Nanopore Technology, SQK-RBK 110.96) was used to sequence the long-read *S. aureus* whole genome on R9.4.1 MinION flowcells (Oxford Nanopore Technology), using 50 ng of the isolated DNA. Base-calling was conducted with Guppy v 6.2.11 in super accuracy mode, using the ’dna_r9.4.1_450bps_sup.cfg’ configuration (Oxford Nanopore Technology). The short-read sequencing was done at a commercial sequencing facility (SA Pathology, Adelaide, SA, Australia) as previously described by Shaghayegh et al. (Shaghayegh et al., 2023). Short-read sequencing was carried out on the Illumina platform, using the Illumina NextSeq 550 (Illumina Inc, San Diego, USA) and NextSeq 500/550 Mid-Output kit v2.5 (Illumina Inc., FC-131-1024). To prepare for short-read sequencing, the genomic DNA was isolated using the NucleoSpin Microbial DNA kit (Machery-Nagel GmbH and Co.KG, 740235.50, Duren, Germany). The sequencing libraries were prepared using a modified protocol for the Nextera XT DNA library preparation kit (Illumina Inc. FC-131-1024). The genomic DNA was fragmented, after which a low-cycle PCR reaction was used to amplify the Nextera XT indices to the DNA fragments. One hundred fifty bp reads were obtained by sequencing after manual purification and normalisation of the amplicon library.

### Bioinformatics

#### Chromosome assembly

We created complete chromosomal assemblies of *S. aureus* using the custom Snakemake pipeline (Molder et al., 2021) that can be accessed via https://github.com/gbouras13/hybracter and a Snaketool (Roach et al., 2022) powered command line tool called hybracter (Bouras, hybracter). Briefly, the long reads were reduced to 250 Mbp for each sample using Rasusa (M. Hall, 2022). Adapters and barcodes were removed using Porechop (Ryan R. Wick, Porechop), short reads were filtered and trimmed with low-quality regions, and adapters were removed using fastp (Chen, Zhou, Chen, & Gu, 2018). Long-read-only assemblies were created using Flye v2.9.1 with the option “--nano-hq.” (Kolmogorov, Yuan, Lin, & Pevzner, 2019). Assemblies, including contigs with a length greater than 2.5 Mb, were kept and denoted as the putative chromosomal. The resulting chromosomes were first polished with long reads using Medaka v1.7.0 (ONT, 2022), then with short reads using Polypolish v0.5.0. (R. R. Wick & Holt, 2022) . After the first round of polishing, the chromosomes were reoriented to start at the *dnaA* gene using the python program called dnaapler (Bouras, dnaapler). Finally, chromosomes were polished for a second time using Polypolish and then with POLCA (Zimin & Salzberg, 2020).

#### Plasmid assembly

Plassembler v 0.1.4 (Bouras, Plassembler) was used to assemble bacterial plasmids from a combination of long and short sequencing reads. Firstly, the short reads are filtered using fastp. The long reads were filtered using nanoFilt (De Coster, D’Hert, Schultz, Cruts, & Van Broeckhoven, 2018) and then assembled using Flye. The largest contig was evaluated to see if the assembly contained more than one contig. If this contig was over 90% of the length of the chromosome size (∼2.5 MB), it was identified as the chromosome. All other contigs were deemed putative plasmid contigs. Both long and short reads were then mapped twice, first to the chromosome and then to the plasmid contigs. For the mapping, minimap2 (H. Li, 2018) was used for the long reads, while BWA-MEM (Heng Li, 2013) was used for the short reads. Reads aligned to the plasmid contigs or not aligned to the chromosome were extracted, combined, and de-duplicated. To produce the final plasmid contigs, these reads were assembled using Unicycler v0.5.0 (R. R. Wick, Judd, Gorrie, & Holt, 2017).

#### Annotation

Chromosome and plasmid assemblies were annotated with Bakta v1.5.0 (Schwengers et al., 2021). The assemblies were typed according to multi-locus sequence typing (MLST) using the program MLST (Seemann, mlst) and assigned to clonal complexes of PubMLST (Jolley, Bray, & Maiden, 2018). Variable-length-k-mer clusters (VLKCs) were used to query the assemblies with k-mer lengths ranging from 13 to 28 and a sketch size of 9984 using the pp-sketchlib tool (Lees et al., 2019). The VLKCs were assigned to the pre-built PopPUNK Staphopia database of 103 clusters for phylogenetic analysis (Petit & Read, 2018). The phylogenetic tree was visualised using the ggtree R package (Xu et al., 2022).

#### Chromosome analysis

The presence or absence of resistance and virulence genes in the genome of the CIs was determined by screening contigs using ABRicate v1.0.1 (Seemann, Abricate) against the Comprehensive Antibiotic Resistance Database (CRAD) (Jia et al., 2017) and the Virulence Factor Database (VFDB) (Liu, Zheng, Jin, Chen, & Yang, 2019).

Genome-wide association analysis was done by first creating a pangenome of the 34 T0 isolates with panaroo v1.3.2 (Tonkin-Hill et al., 2020) and then testing the significance of each gene with Scoary v1.6.16 using default parameters (Brynildsrud, Bohlin, Scheffer, & Eldholm, 2016). All following paired *S. aureus* genomic analysis was conducted using a Snakemake pipeline. Firstly, small variants, such as single nucleotide variants (SNVs) and small insertions and deletions, were called using Snippy v 4.6.0. (Seemann, 2015), with the raw FASTQ short reads from the Timepoint T1 isolates were compared against the corresponding GenBank file of the assembled Timepoint T0 isolate for each CIs pair. All larger structural differences were called using two methods: Nucdiff v2.0.3 (Khelik, Lagesen, Sandve, Rognes, & Nederbragt, 2017) and Sniffles v2.0.7 (Sedlazeck et al., 2018). For Nucdiff, chromosome assembly of the T0 isolate was compared against the corresponding T1 isolate. For, Sniffles, all T1 isolate long reads were first aligned to the T0 isolate genome using minimap2 v 2.24 (H. Li, 2018) specifying ’-ax map-ont’ parameters. The resulting BAM was used as input for Sniffles.

The large structural variant CIs pairs of subject 420 and 4875 were manually annotated by mapping all timepoint T1 long reads to the T0 assembly using minimap2 v 2.24 specifying ’-ax map-ont’, followed by sorting the resulting BAM file using samtools (Danecek et al., 2021). Structural deletions were visualised in R using the gggenomes, and the long-read pile-up was visualised using the Gviz packages (Ankenbrand, 2022; Hahne & Ivanek, 2016).

#### Plasmid analysis

For each putative plasmid contig derived from the output of Plassembler, Mobtyper v1.4.9 (Robertson & Nash, 2018) was run to determine each plasmid’s predicted mobility and replicon marker. The minhash (’Mash’) distance was calculated between each pair of plasmids using mash v2.3 (Ondov et al., 2016). A plasmid pangenome was created using panaroo v1.3.2. To determine shared plasmid genes using the ‘gene_presence_absence. Rtab’ output, the Jaccard index based on gene presence and absence, was calculated between each plasmid pair. Following the analysis by Hawkey et al., plasmids were empirically determined to be the same plasmid using thresholds of Mash similarity > 0.98 and Jaccard index > 0.7 (Hawkey et al., 2022).

Additionally, plasmids were determined to be beta-lactamase-carrying if they carried the *blaZ*, *blaI* and *blaR1* gene operon. All plasmid-copy numbers were obtained using Plassembler v0.1.4.

### Relatedness of isolate pairs

A two-step approach was used to classify isolate pairs as either closely related ’same strain’ or not closely related ’different strain’. Firstly, the Sequence Types obtained from MLST and clusters generated by PopPUNK were compared between each isolate pair. If either of these metrics differed in the CI pair, they were considered to belong to different strains. The second step involved analysing the number of small variants outputted by Snippy. Isolate pairs with 100 or fewer single nucleotide variants (SNVs) between the first and second timepoint were classified as the same strain.

### Planktonic Antibiotic Susceptibility

Susceptibility testing followed Clinical and Laboratory Standards Institute (CLSI) guidelines (CLSI, 2020). Seven antibiotics were chosen for susceptibility testing according to their common use in medical practice. These were: amoxicillin in combination with clavulanic acid (augmentin), clarithromycin, clindamycin, doxycycline, erythromycin, gentamicin, and mupirocin (Sigma-Aldrich, St. Louis, USA). Minimum Inhibitory Concentrations (MICs) were obtained for the planktonic form of all isolates, utilising the microbroth dilution assay (Wiegand, Hilpert, & Hancock, 2008). The antibiotics were tested at 0.06–32 mg/L dilution range. The assay was repeated at least twice per CI. The MIC50, MIC90 and antibiotic non-susceptibility proportions were calculated adopting the susceptibility breakpoints published by the CLSI.

### Biofilm Antibiotic Tolerance

The biofilm tolerance assay was based on a 96-well plate adapting the procedures used by Mah et al. (Mah, 2014). Each isolate was exposed to the same antibiotics used for the planktonic antibiotic susceptibility testing. The concentrations ranged from 1.25–640 mg/L. In brief, the CIs were cultured on Mueller-Hinton agar (Sigma-Aldrich). Then, single colonies of *S. aureus* were suspended in 0.9 % saline to a turbidity reading of 0.5 McFarland Units (MFU). The 0.5 MFU bacterial suspension was diluted 100-fold in Mueller-Hinton broth to achieve a 5 × 10 ^5^ CFU/ml before inoculation in a 96-well plate (200 μL). Plates underwent a 48-hour incubation at 37°C with sheer force on a rotating plate set at 70 rpm (3D Gyratory Mixer, Ratek Instruments, Australia). Following the incubation, the supernatants were gently aspirated with a minimum agitation of the biofilms. These biofilms were then exposed to different antibiotics in serial diluted (200 μL) Mueller-Hinton broth for 24 hours. After incubation with antibiotics, the supernatants were aspirated gently, and non-adherent planktonic bacteria were removed by gently washing with sterile phosphate-buffered saline (PBS). Subsequently, the biofilm tolerance was assayed using a resazurin viability method, alamarBlue Cell Viability Reagent (Thermo Fisher Scientific, DAL1025), as per the manufacturer’s instruction (Pettit et al., 2005). The assay was repeated twice per CI with two replicates.

### Biofilm Biomass Assay

To quantify the total biofilm biomass, the Crystal Violet (CV) staining assay was used (Stiefel et al., 2016). Inoculated 96-well plates underwent a 48-hour incubation at 37°C on a rotating plate set at 70 rpm to induce biofilm formations. Following the incubation, the planktonic cells were removed by gently aspirating the supernatants and washing the wells twice with PBS. Subsequently, 200 μL of 0.1 % CV (Sigma-Aldrich, C6158) solution was added for 15 minutes. After washing the wells three times with sterile water and air-drying, the fixed CV was solubilised by adding 200 μL 30% acetic acid and shaking for one hour at room temperature. The absorbance was obtained at 595 nm with a FLUOstar Omega microplate reader (BMG Labtech, Ortenberg, Germany). The assay was repeated twice per strain, with six technical replicates.

### Statistics

We used a generalised linear mixed model (GLMM) to analyse the antibiotic tolerance data. To assess the significance of each variable, a backwards stepwise regression approach using the log-likelihood ratio test was used to remove insignificant variables. The threshold of significance was set at a p-value<0.05. All analysis was performed with R v4.2.0. (R Core Team, 2017).

### Data availability

The assembled chromosomes and plasmids, raw short and raw long read FASTQs, are accessible on the Sequence Read Archive (SRA) under the project code: PRJNA914892. The complete list of biosample accession numbers for each sample can be found in supplementary table 1.

### Code Availability

All code used to generate all analyses & figures used in this manuscript can be found at https://github.com/gbouras13/CRS_Saureus_Evolutionary_Landscape.

## Results

### Clinical characteristics

Thirty-four *S. aureus* sequential pairs (68 clinical isolates from which 34 first timepoint (T0) and 34 second (T1) isolates) were included in this study, isolated from 34 subjects. The mean time between paired *S. aureus* CI collection was 18 months (range 6-52). Most subjects were classified as CRSwNP (85%) and having asthma (56%). The clinical characteristics of the subjects are summarised in Table S2.

### *S. aureus* strains persist within the sinonasal cavities in 41% of cases

The MLST analysis revealed a total of seven clonal complexes (CCs), including CC1 (n=5, 7.3%), CC5 (n=4, 5.8%), CC8 (n=2, 2.9%), CC15 (n=5, 7.3%), CC22 (n=5, 7.3%), CC30 (n=9, 13.2%), and CC45 (n=14, 20.5%). A total of 24/68 CIs (35.2%) were not assigned to any CC (Fig. 1A). The analysis of the PopPUNK variable-length-k-mer clusters (VLKCs) identified a total of 16 clusters. Of the 34 isolate pairs, 18 (52.9%) pairs belonged to different CCs or VLKCs, indicating that they were not closely related isolates and were classified as ’different strain’ pairs.

**Figure 1.**
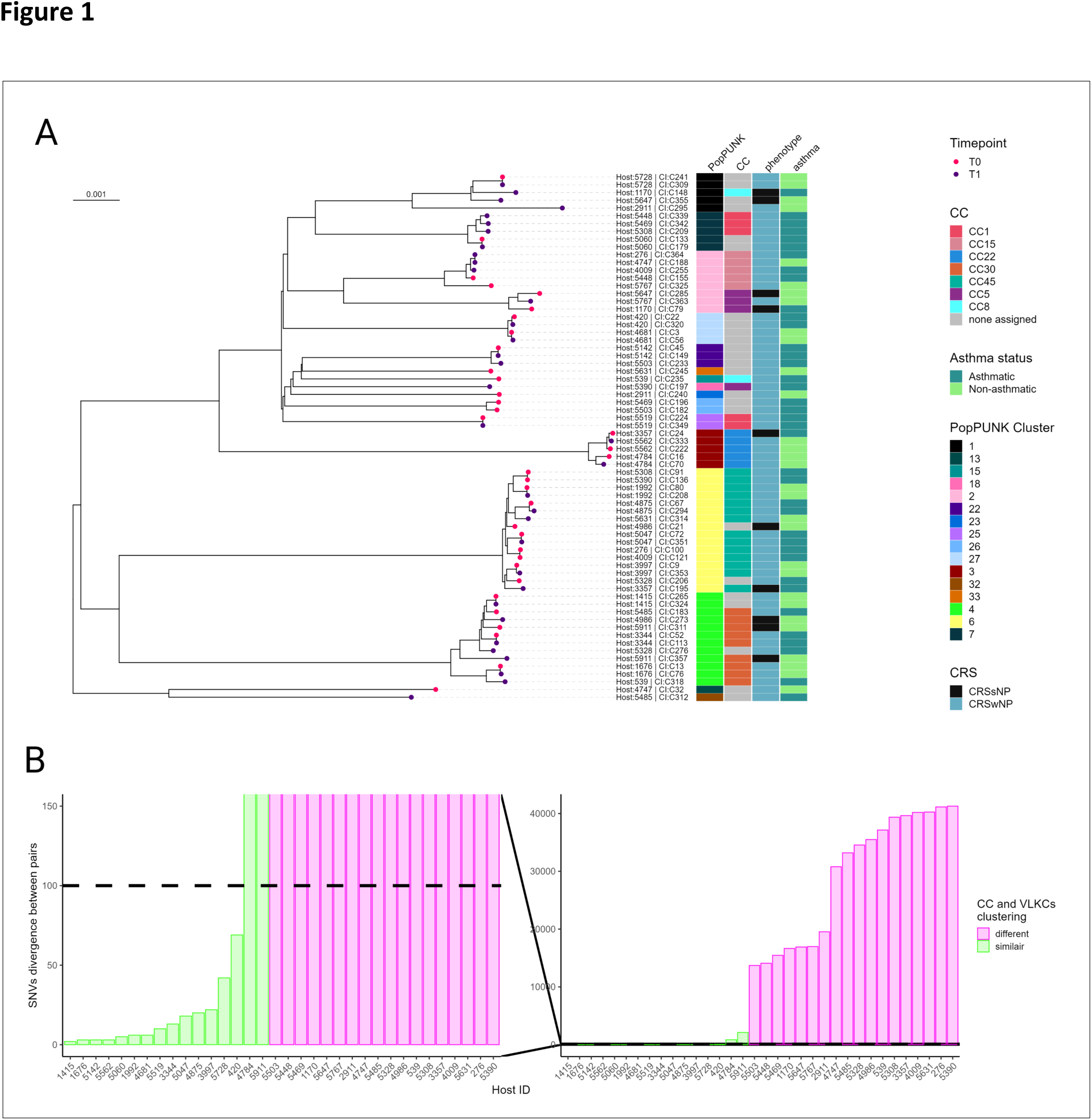
Genome-based classification of *Staphylococcus aureus* clinical isolates. (A) A variable-length-k-mer cluster (VLKC) midpoint rooted tree of 68 *S. aureus* genomes collected from 34 subjects with chronic rhinosinusitis (2 samples per subject) based on PopPUNK analysis. The branch tip colours represent the collection timepoint (T0= first, T1=later timepoint). The PopPUNK cluster, clonal complex (CC), CRS phenotype, and asthma status are indicated by colour on the right side. The branch labels show the corresponding host ID and the CI number. (B) A histogram depicting the distribution of pairwise single-nucleotide variant (SNV) divergence in the core genome for all clinical isolate pairs (n=34), with colours indicating CC and VLKC similarity. The horizontal line indicates the SNV threshold used to classify pairs as either “same strain” or “different strain”.

The relatedness was investigated for the 16 remaining pairs by analysing the number of SNVs in shared genes. The variants ranged from 2-2123, with 14/16 pairs having 69 or fewer SNVs and 2/16 pairs having more than 100 SNVs. Figure 1B shows the multimodal distribution of SNVs between all CI pairs and reflects the classification of pairs into different and same strain groupings. The two pairs with the same CC and VLKC groups and more than 100 SNVs divergence (host 4784, 846 SNVs; host 5911, 2123 SNVs) were classified as ’different strain’ pairs due to the large number of SNVs and structural variations between pairs. Namely, the CI pair from host 4784 had 100 structural variations between T0 and T1 according to Sniffles, with the addition of a plasmid in the T1 isolate. Similarly, the CI pair from host 5911 had 143 structural variations between them. Accordingly, 14/34 pairs (41%) were classified as unambiguously in the ’same strain’ group of persistent isolates, whilst 20/34 pairs (59%) were classified as being part of the ’different strain’ group, where the subject had been colonised or infected by a different strain over time (Table S3). No genomic clustering was observed based on the order of CI collection of the pairs and the host’s CRS phenotype or asthma status (Fig. 1A).

### Chromosomally encoded antimicrobial resistance genes and virulence factors are widespread in *S. aureus* sinonasal isolates

Chromosomally encoded antimicrobial resistance (AMR) genes in the *S. aureus* isolates were assessed using the CARD database, revealing a range of 8-21 genes AMR per isolate. Most isolates (67/68) contained 8-13 AMR genes, including *arlR*, *arLR*, *arlS*, *lmrS*, *mepA*, *mepR*, *mgrA*, *norA* and *tet(38),* which were identified in all CIs (Fig. S1). Only one isolate (Host:2911, CI: C295) contained more than 13 AMR genes. Among the 40 CIs classified as being different strain pairs, the *blaZ* beta-lactamase gene was present in 22 of them. Notably, the prevalence of chromosomal *blaZ*-positive isolates increased from 9/20 (45%) in the first timepoint different strain group to 13/20 (65%) in the second timepoint different strain group, indicating a potential selective pressure for beta-lactamase-resistant isolates in the population. Remarkably, none of the isolates in the second timepoint of the different strain group contained the *ermC* gene, whereas, in the first timepoint, three isolates were found to carry multiple copies.

In the same strain group isolates, 11/28 (39.2%) were positive for a chromosomally encoded *BlaZ* beta-lactamase gene. Only one of the same strain pairs gained a chromosomally encoded *BlaZ* gene at the second timepoint (Fig. S1).

The presence of chromosomally encoded virulence factor genes in the *S. aureus* isolates was assessed using the VFDB database, revealing a range of 45-72 (median 57) genes per isolate (Fig. S2). Notably, all CIs contained the serine protease operon sspABC, also known as V8 protease, which has been previously associated with allergic sensitisation to *S. aureus* (Krysko, Teufelberger, Van Nevel, Krysko, & Bachert, 2019). Additionally, all isolates had immune evasion-associated factors such as the immunoglobulin-binding protein *sbi*, *adsA*, *lip*, *hly/hla*, *hlgAB*, *hld*, and *geh*. The *isdABCDEFG* operon was present in 67 out of 68 isolates. The *icaABCD* operon, associated with biofilm production, was present in all isolates, but interestingly, two isolates lacked the *icaR* (repressor) gene.

Moreover, 61 and 66 isolates contained the *sak* and *scn* virulence factors, respectively, which are prophage encoded (Nepal et al., 2021). Notably, the prevalence of immune evasion factors *chp* (9/20 vs. 18/20) and *sdrE* (9/20 vs. 15/20) increased in the second timepoint different strain group. In contrast, the carriage of *sdrC* (13/20 vs. 7/20) decreased in the second timepoint of different strain group (Fig. S2). No remarkable alterations were observed in the acquisition or loss of virulence factors between the initial and subsequent timepoints of the same strain group.

### Analysis of gene content in *S. aureus* isolates suggests a reduced virulence profile for incoming isolates

To investigate whether gene presence or absence was linked to persistence, a microbial gene presence-absence analysis was performed on the 34 Timepoint T0 isolates using Scoary. No statistically significant differences in gene content were found between the same and different strain isolates at T0 (BH p.adj > 0.05). However, the *chp* gene, involved in chemotaxis inhibition, was less prevalent in the persister same strain group (8/14 same strain vs. 18/20 different strain).

A subsequent microbial gene presence-absence analysis was conducted on the 40 different strain isolates to examine whether the gene content of the second timepoint T1 isolates differed from that of the T0 isolates. Although no statistically significant differences were observed, there was a clear trend for incoming T1 isolates to have fewer virulence factors than the T0 isolates they replaced, such as staphylococcal enterotoxins M, U, I, N, and G (present in 8/20 T1 isolates compared to 18/20 T0 isolates) and *chp* (9/20 T1 vs 18/20 T0 isolates).

### Same strain SNV prevalence reveals heterogeneous host adaption

222 SNVs were observed across the 14 same-strain isolates, ranging from 2-69 per isolate. 8/14 pairs had less than 10 SNVs. Of these 222 SNVs, 148 were in putative coding sequences (CDS), 44 were synonymous SNPs, 3 were in-frame variants, 9 were frameshift variants, 4 were stop-gained variants, and the remaining 88 were missense- SNVs. Only three genes contained SNVs in more than one isolate, namely the ribosomal protein *rpsJ,* the transcription termination factor *clpC*, and the protease ATP-binding subunit *clpX*. Interestingly, MSCRAMM genes commonly harboured SNVs across the same strain pairs, with 11 SNVs occurring in 9 distinct MSCRAMM genes in 6 distinct isolates, of which 6/11 mutations were synonymous. These include variants in the *sdrC* adhesin, fibronectin-binding proteins A and B (*fnbA* and *fnbB*), surface protein G (*sasG*) and iron-regulated surface-determining proteins *isdD*, *isdE* and *isdF*. Other adhesion genes, including Staphylocoagulase *coa*, extracellular adherence protein *eap*/map, and the extracellular matrix binding protein *EbhA* also had SNVs across isolates.

### Structural variants in same strain CI pairs involve prophages, insertion sequences, MSCRAMM and AMR Genes & are not correlated with the number of SNVs

We detected a total of 37 structural variants (SV) among the 14 same strain isolates, ranging in size from small collapsed duplications (<10bp) to the acquisition of a 43793 bp *hlb*-disrupting Sa3int prophage in a single isolate pair. Only 10 SV were larger than 100bp, and all were found in 4/14 CI pairs, with one strain having 5 SV> 100bp. Notably, no relationship was observed between the number of SNVs and structural variations. The CI pair from host 5562, which had the second-lowest number of SNVs (3), had 5 SV, while 10 strains did not have any SV> 100bp, including 4 strains with > 10 SNVs. In addition, 5 insertion sequence (IS) insertions were identified in 3 distinct strains, one of which disrupted the *agr* locus (Table 2).

**Table 2.** SNV and SV count between same strain isolate pairs.

| Host ID CI pair | SNV count (Snippy) | SV count (Nucdiff) | SV count (Sniffles) |
| --- | --- | --- | --- |
| 1415 | 2 | 1 | 2 |
| 1676 | 3 | 1 | 1 |
| 1992 | 6 | 4 | 3 |
| 3344 | 13 | 0 | 1 |
| 3997 | 22 | 0 | 2 |
| 420 | 69 | 5 | 5 |
| 4681 | 6 | 2 | 1 |
| 4875 | 20 | 7 | 5 |
| 5047 | 18 | 1 | 2 |
| 5060 | 5 | 0 | 0 |
| 5142 | 3 | 0 | 0 |
| 5519 | 10 | 2 | 1 |
| 5562 | 3 | 11 | 8 |
| 5728 | 42 | 3 | 4 |

SNV, Single nucleotide variant; SV, Structural Variant

Interestingly, between the same strain CI pairs obtained from host subject 420, there was a 4638 bp deletion between T0 and T1. This deletion encompassed the cell-wall spanning region, the transmembrane region, and the cytoplasmic domain of the MSCRAMM serine-repeat *sdrC* gene, along with the signal sequence, ligand binding domain and repeat regions in the neighbouring serine-repeat *sdrD* gene as depicted in the coverage and pile-up plot shown in Figure 2B. This was leading to the recombination of the cell-wall spanning region, the transmembrane region and the cytoplasmic domain from the *sdrD* gene with the signal sequence, ligand binding domain, and repeat regions of the sdrC gene (Fig. 2A). Additionally, in this CI pair, the fibrinogen-binding adhesin *SdrG* had a tandem duplication, and there was a tandem duplication in the extracellular adherence protein *Eap/Map* over time.

**Figure 2.**
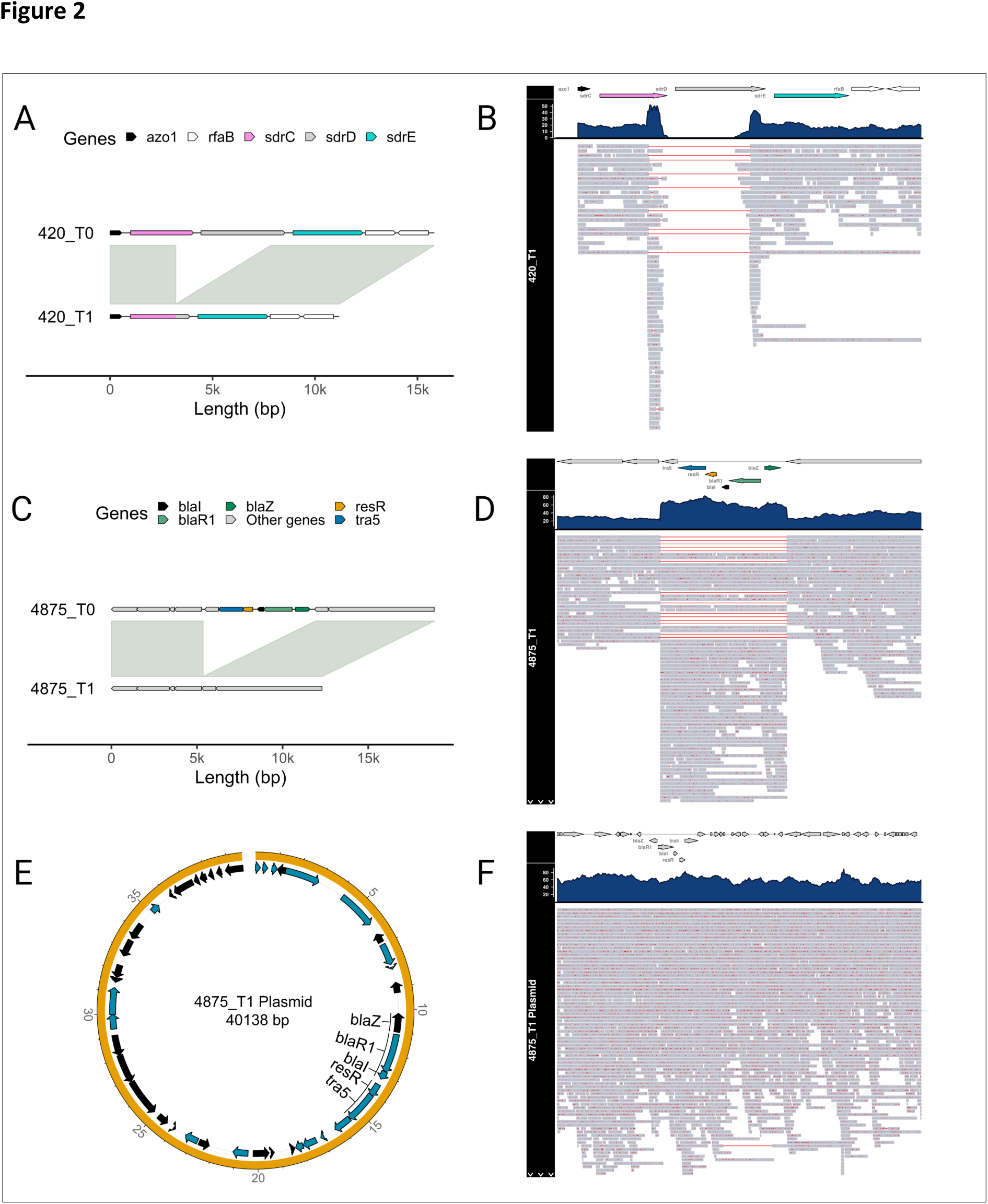
Structural variants identified between same strain longitudinal pairs. (A) Alignment of the *sdrCDE* locus between two isolates from the same host (420) at different timepoints. Genes are highlighted in different colours, and synteny and sequence similarity are indicated by grey fills connecting the chromosomes. The genomes of the first and second timepoint isolates are shown on top and bottom, respectively. (B) Coverage and pile-up plot of aligned long-reads of the second timepoint isolate of host 420 against the *sdrCDE* locus of the first timepoint isolate, red indicates deleted regions in the reads. (C) Alignment of the *β-lactamase* locus between two isolates from the same host (4875) at different timepoints. (D) Coverage and pile-up plot of aligned long-reads of the second timepoint isolate of host 4875 against the *β- lactamase* locus of the first timepoint isolate, red indicates deleted regions in the reads. (E) Circular plot of the acquired plasmid of the second timepoint isolate of host 4875. (F) Coverage and pile-up plot of aligned long-reads of the acquired plasmid of the second timepoint isolate of host 4875, red indicates deleted regions in the reads.

Another noteworthy observation was the identification of a large SV event between the same strain pair from host 4875. A transposon carrying the blaZ locus was lost in the second timepoints isolate (Fig. 2C). Specifically, the second timepoint isolate showed a loss of a transposon carrying the *blaZ* locus in the chromosome while simultaneously acquiring a plasmid containing the same locus (Fig. 2C and E). The coverage and pile-up plot shown in Figure 2D revealed that the coverage of the *blaZ* locus contained by the plasmid was higher compared to that of the chromosome, likely due to the high plasmid copy number (Fig. 2F). These findings highlight the dynamic nature of virulence and AMR genes that occur in persistent *S. aureus* isolates.

### Plasmid carriage is common, and plasmids often encode beta-lactamase resistance genes

Hybrid long and short-read sequencing allowed us to analyse the plasmid content of these isolates and probe their change over time. Fifty-three plasmid contigs were assembled from 41/68 isolates, while the remaining 27 isolates did not carry any plasmids. 43/53 plasmid contigs were determined to be complete and circularised, while the other 10 were putative incomplete contigs. The analysis of the plasmids detected in the 68 CIs revealed a bimodal distribution of mash distances between each plasmid contig (1^st^ quantile: 0.0, Median: 0.85, 3^rd^ quantile: 0.94), indicating that approximately 50 plasmids had a high level of similarity (Fig. 3). Twenty plasmid contigs were identified in the 28 ’same strain’ isolate pairs, of which 16 (8 at each timepoint) were present in both timepoints. Plasmid gain was observed between 2 isolate pairs (subjects 3997 and 4875), where the second timepoint isolates C353 and C294 gained 1 and 2 plasmids, respectively. In contrast, plasmid loss was observed in one same strain isolate pair (5047), where the second timepoint isolate C351 lost one plasmid over time. 27/53 plasmid contigs from 26 distinct isolates carried the beta-lactamase gene *blaZ*. Of these 27 plasmids, 17/27 were deemed closely related enough to be analysed as the same plasmid based on the empirical thresholds outlined in the methods (Fig. S3). Two additional antimicrobial resistance genes were identified in plasmids: erythromycin resistance gene *ErmC* (encoded on a 2473bp plasmids common to 3 strains) and the quaternary ammonium compound resistance gene *qacA*, found on 1 20560bp plasmid.

**Figure 3.**
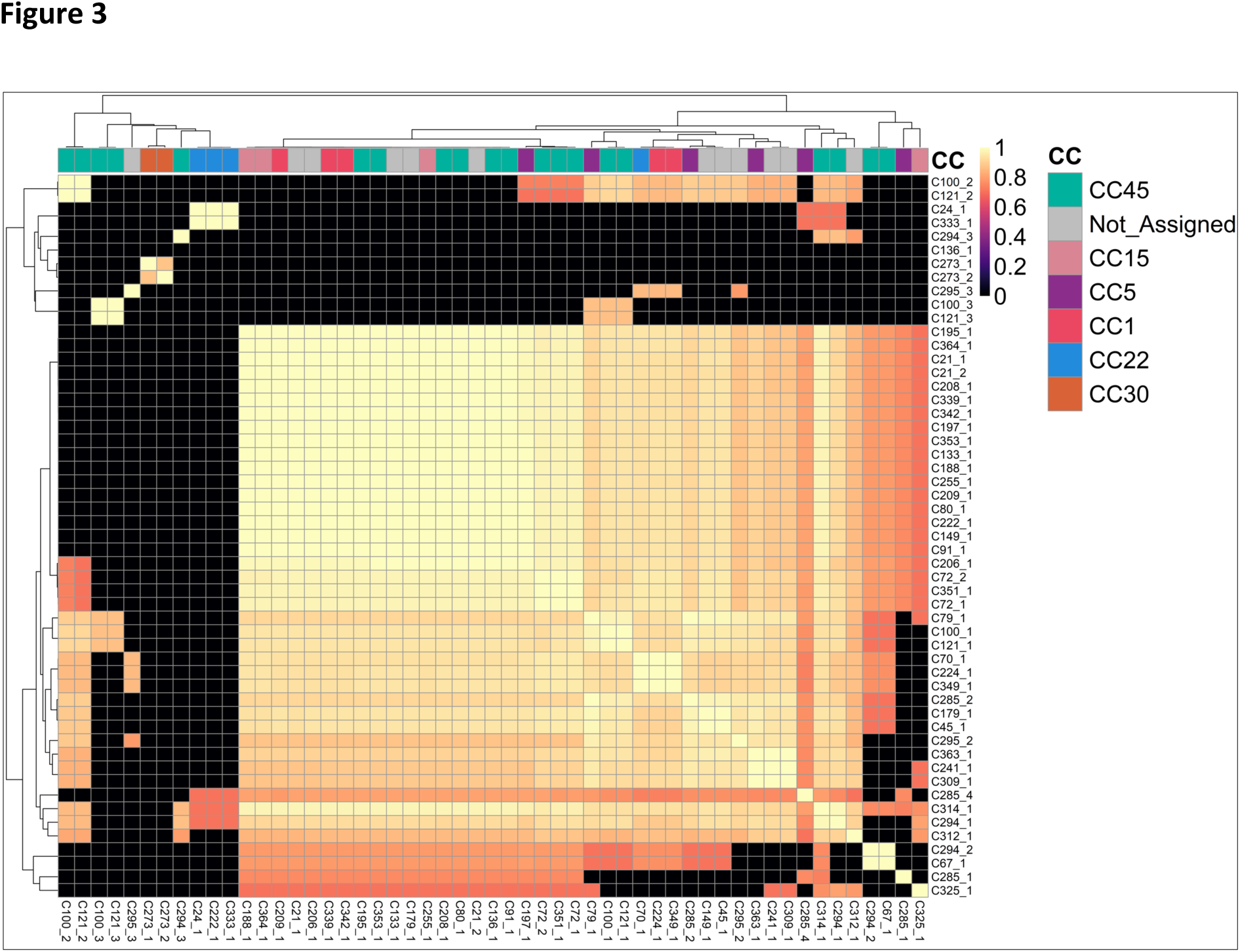
Heatmap displaying the minhash (Mash) distances between the 53 plasmids identified in the 68 CIs. The distances were calculated using mash v2.3 and are represented by a colour gradient. The clonal complex of the CI from which the plasmid was recovered is indicated at the top of the heatmap.

The number of beta-lactamase encoding plasmids increased from 11 at T0 to 16 at T1, with two same strain isolates acquiring beta-lactamase resistance plasmids. In contrast, three different strain isolates present at T1 replaced isolates at T0 that did not carry a beta-lactamase plasmid, indicating a selection pressure to gain beta-lactamase resistance.

### Plasmid copy numbers increase with time in the same strain group

We found a moderate positive correlation (Spearman’s correlation coefficient R=0.63) between plasmid copy numbers estimated using long and short-read methods (Fig. S4A). The median plasmid copy number estimation was 1.63 times higher in the long-read dataset than in the short-read dataset. Beta-lactamase-carrying plasmids exhibited an even more noticeable difference in copy number estimation between techniques, with a median 2.29 times higher copy number estimate in the long-read dataset. The long-read dataset did not capture four plasmid contigs; however, these were all incomplete (Fig. S4B).

We further investigated the stability of plasmid copy numbers in the ’same strain’ group, focusing on the eight conserved plasmids. We observed a significant increase in the copy number of the conserved plasmids over time (p<0.05) (Fig. 4), with 4 of the eight conserved isolates being *blaZ* positive. However, we observed no significant difference between the plasmid copy number and timepoint when we examined the short-read data.

**Figure 4.**
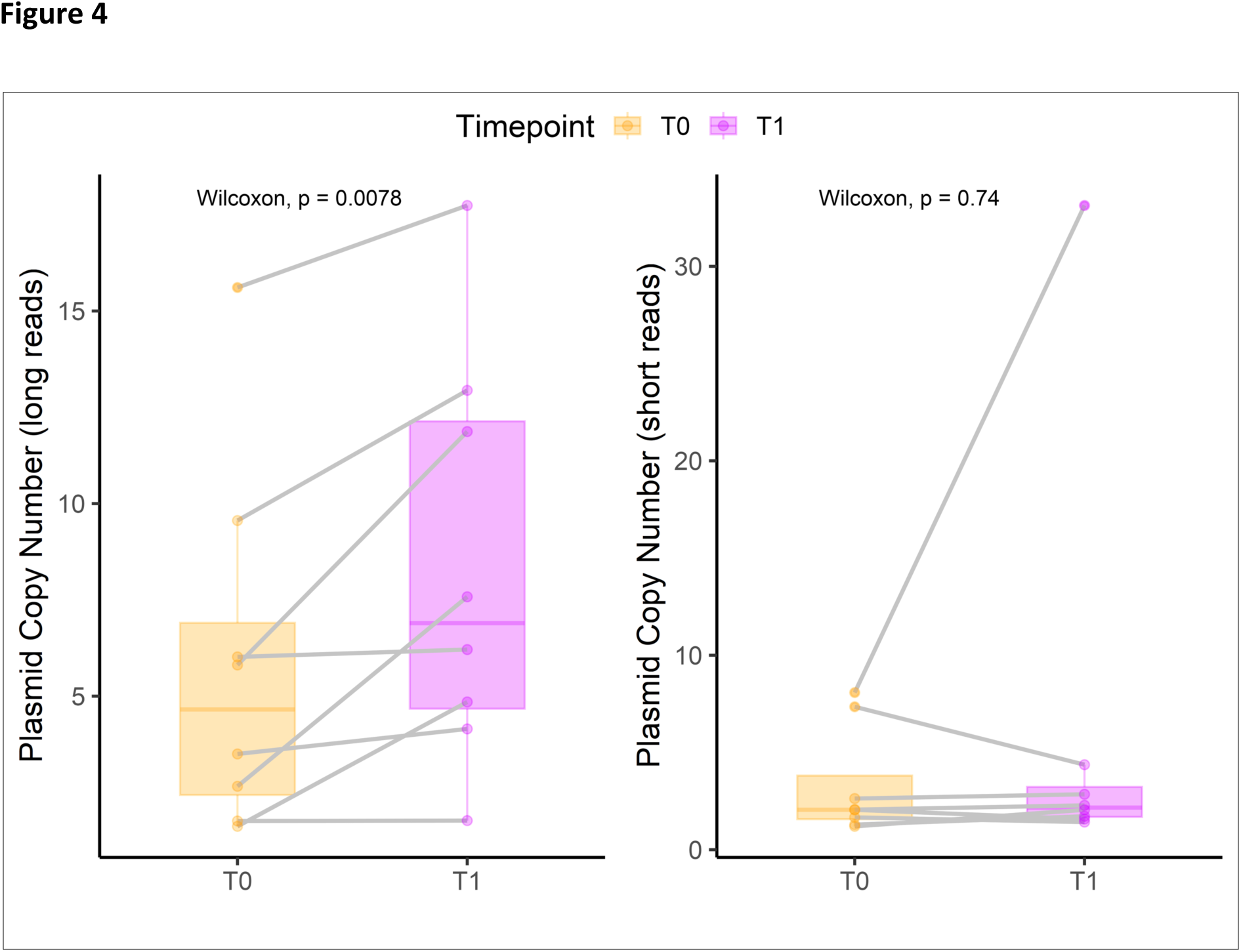
Copy numbers of the conserved plasmids in the ’same strain’ group (n=8) for short-read and long-read data. The colour indicates timepoints, and the grey line indicates paired conserved plasmids. The Wilcoxon signed-rank test compared the copy numbers between the two timepoints, with p<0.05 considered significant.

### Planktonic antibiotic susceptibility remains stable over time

The antibiotic susceptibility of all CIs was tested (n=68). Mupirocin appeared to be the most potent, with 85.2% and 67.65% of MIC values below the lowest concentration tested (0.06 mg/L) for the first and second timepoints CIs, respectively. In contrast, erythromycin and clarithromycin had lower susceptibility rates, with over 22% of CIs being resistant to each antibiotic (Table S4). Overall, doxycycline was highly effective at both timepoints, with 97% of CIs being susceptible. When comparing the first and second timepoints CIs, there was no significant difference in the proportion of resistance between the CI pairs classified in the different or same strain group (Fig. S5).

### Biofilm antibiotic tolerance increases over time in persistent *S. aureus* strains

Next, we investigated the antibiotic tolerance of biofilms for all isolates. The viability results after antibiotic treatment were analysed using a GLMM. The model included the following variables: timepoint, antibiotic, antibiotic concentration, and same strain-relatedness classification. The summary statistics of the GLMM results for all effects are provided in table 1. The biofilm viability data showed high variability in antibiotic tolerance between CIs and antibiotics. Although all antibiotics significantly reduced the biofilm viability (p<0.001), their dose-response relationships varied. Except for doxycycline, all antibiotics reached a plateau in their antibiofilm effects at 5 mg/L, reducing biofilm viability by approximately 35%, and did not eradicate biofilms at the highest concentration (640 mg/L). Notably, mupirocin at the lowest concentration of 1.25 mg/L showed a reduction of over 50% in biofilm viability, despite not eradicating the biofilms at 640 mg/L (Fig. 5A).

**Table 1.** General linear mixed-effect model of biofilm viability.

|  | fluorescence |  |  |  |  |
| --- | --- | --- | --- | --- | --- |
| Predictors | Estimates | std. Error | std. Beta | standardised std. Error | p |
| (Intercept) | 9521.99 | 468.05 | 1.47 | 0.16 | *** |
| group [T1] | 80.34 | 69.10 | 0.03 | 0.02 | NS |
| same strain [Yes] | -732.07 | 642.38 | -0.26 | 0.23 | NS |
| concentration [1.25] | -2151.71 | 247.48 | -0.76 | 0.09 | *** |
| concentration [2.5] | -2993.73 | 247.48 | -1.05 | 0.09 | *** |
| concentration [5] | -3375.66 | 247.48 | -1.19 | 0.09 | *** |
| concentration [10] | -3468.68 | 247.48 | -1.22 | 0.09 | *** |
| concentration [20] | -3504.01 | 247.48 | -1.23 | 0.09 | *** |
| concentration [40] | -3664.02 | 247.48 | -1.29 | 0.09 | *** |
| concentration [80] | -3865.20 | 247.48 | -1.36 | 0.09 | *** |
| concentration [160] | -3876.89 | 247.48 | -1.37 | 0.09 | *** |
| concentration [320] | -4123.81 | 247.48 | -1.45 | 0.09 | *** |
| concentration [640] | -3970.43 | 247.48 | -1.40 | 0.09 | *** |
| antibiotic[clarithromycin] | -913.07 | 99.85 | -0.32 | 0.04 | *** |
| antibiotic [clindamycin] | -940.96 | 99.85 | -0.33 | 0.04 | *** |
| antibiotic [doxycycline] | -2226.13 | 99.85 | -0.78 | 0.04 | *** |
| antibiotic [erythromycin] | 465.69 | 99.85 | 0.16 | 0.04 | *** |
| antibiotic [gentamicin] | -234.94 | 99.85 | -0.08 | 0.04 | * |
| antibiotic [mupirocin] | -1526.72 | 99.85 | -0.54 | 0.04 | *** |
| group [T1] Å— same strain<br>[Yes] | 1475.91 | 107.68 | 0.52 | 0.04 | *** |
| Random Effects |  |  |  |  |  |
| σ² | 3389981.04 |  |  |  |  |
| τ₀₀ id | 3350609.42 |  |  |  |  |
| N id | 34 |  |  |  |  |
| Observations | 4828 |  |  |  |  |
| Marginal R² / Conditional R² | 0.184 / 0.590 |  |  |  |  |
| * p<0.05 ** p<0.01 *** p<0.001 |  |  |  |  |  |

**Figure 5.**
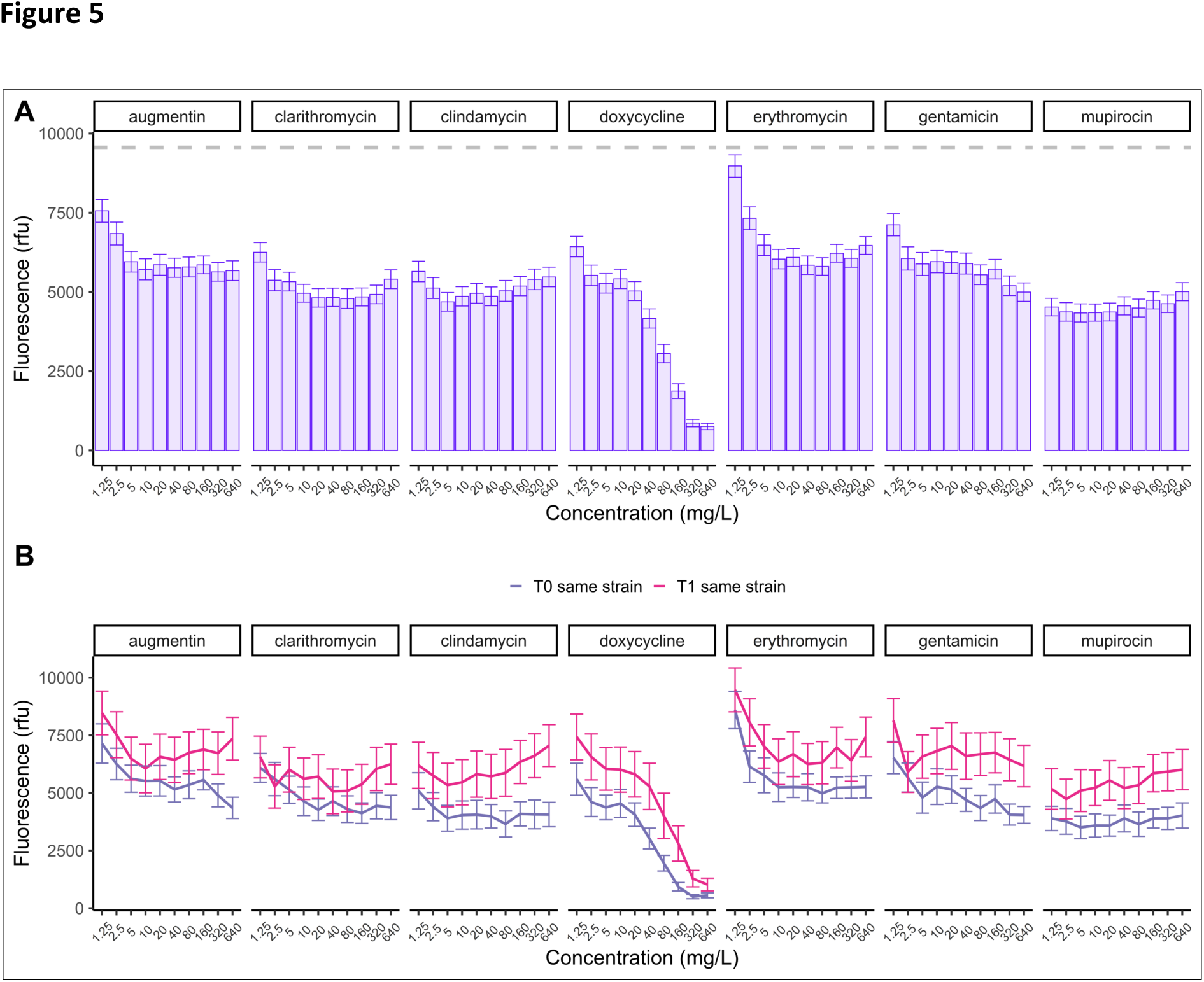
Tolerance of *S. aureus* biofilms to antibiotics. (A) Mean biofilm viability after treatment per antibiotic and concentration in relative fluorescence units (rfu) for all 68 CIs. The grey dashed line represents the mean viability of isolates untreated. (B) Mean biofilm viability of the first and second CIs pairs classified as the same strain after treatment with increasing concentrations of antibiotics. Error bars represent the standard error of the mean (SEM).

Interestingly, we observed a significant increase (p<0.001) in antibiotic tolerance of biofilms over time between the first and second isolates classified as ’same strain’ isolates compared to the first timepoint (Fig. 5B), suggesting that the same strain isolates gained tolerance over time. We then assessed the biofilm biomass using crystal violet staining to investigate the potential relationship between increased antibiotic tolerance and biofilm quantity. We observed a significant increase in the mean biomass of biofilms between the first and second timepoint CIs of the same strain group (paired Wilcoxon signed-rank test, p < 0.05) (Fig. 6), indicating that the increased biofilm tolerance could be due to increased biofilm production by the same strain isolates over time. A similar trend was seen for the biofilm viability results after 48 hours of growth without antibiotic treatment. Specifically, the biofilm fluorescence of the same strain isolates at the first timepoint was significantly lower than that of different strain isolates (p < 0.05). However, the second timepoint of the same strain group showed a significant increase in biofilm production (p < 0.01) over time, resulting in no significant difference in biofilm fluorescence between the second timepoint isolates of the same strain and different strain groups (Fig. S7).

**Figure 6.**
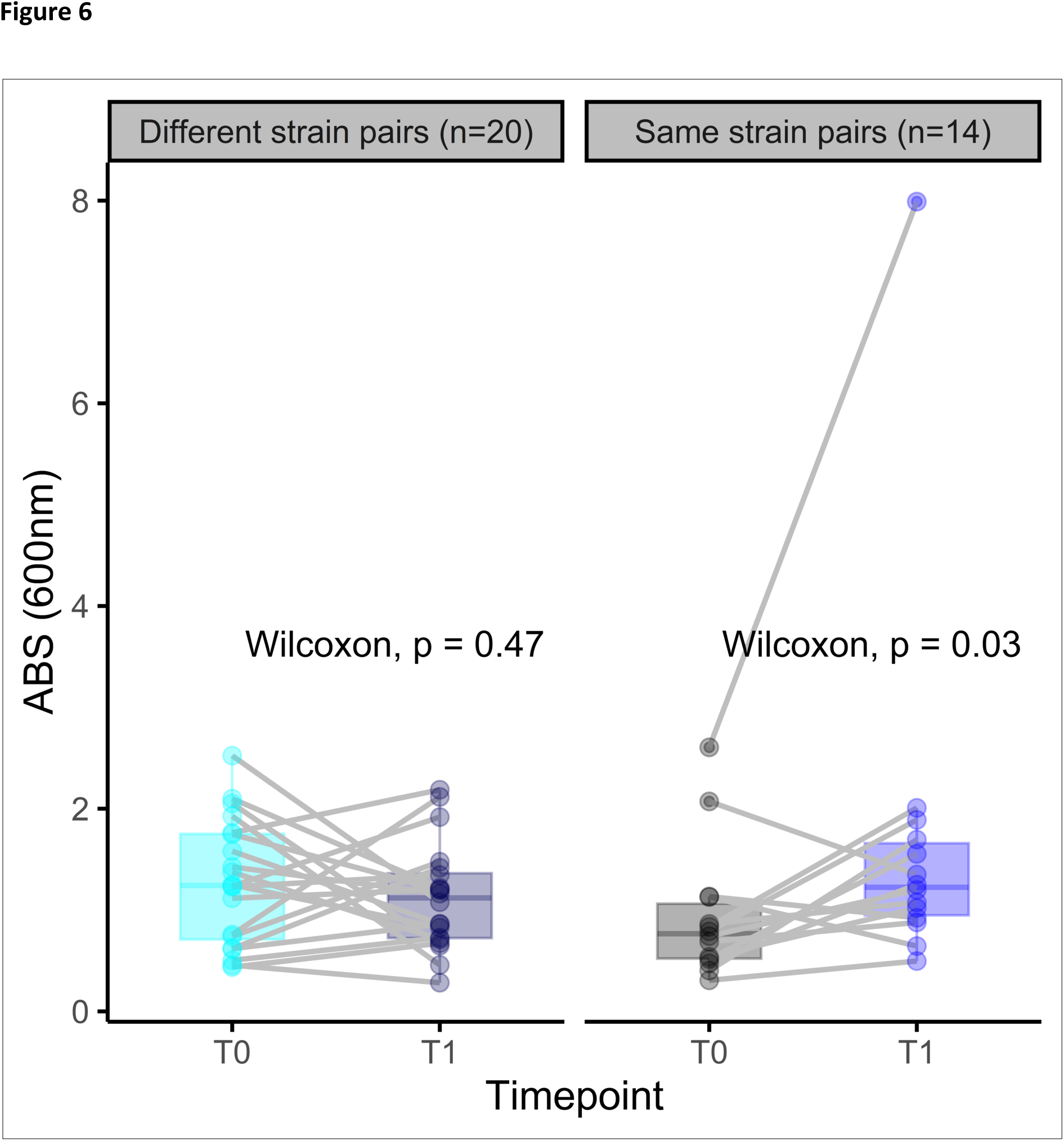
Biomass of *S. aureus* biofilms classified as same strain and different strain using the crystal violet assay. The CIs pairs are connected with a grey line. Paired Wilcoxon test was used to determine the significance between the first and second timepoint.

Next, we investigated the potential correlation between the number of days of antibiotic usage by CRS patients and the increased biofilm antibiotic tolerance. The most commonly prescribed antibiotic was augmentin, but in terms of total exposure time, doxycycline, followed by sinus/nasal saline irrigation mixed with mupirocin and augmentin, had the most extensive antibiotic exposure in all subjects (Fig. S6). CIs classified as the same and different strains had a mean exposure of 16.4 days (±7.97) and 15.7 days (±8.39), respectively. However, we did not find a significant relationship between the total number of days of antibiotic exposure and increased biomass between CIs pairs (Spearman correlation coefficient = -0.11, p = 0.58).

## Discussion

The current study aimed to investigate the persistence of *S. aureus* in the nasal cavity of chronically colonised CRS patients and the related genomic and phenotypic changes over time in a set of longitudinal collections of *S. aureus* CIs. Our hybrid long and short- read sequencing approach allowed us to assemble near-perfect complete genomes and conduct detailed longitudinal genomic analysis. While our study did not identify a specific gene or gene cluster that explains *S. aureus* persistence, persister isolates often show changes in mobile genetic elements such as plasmids, prophages and insertion sequences, indicating a role of the ’mobilome’ in promoting persistence. The genomic adaptation of persister isolates was episode-specific, suggesting that each colonisation event may select different adaptations that enable the survival of *S. aureus* in each host. Additionally, the increase in biofilm tolerance to antibiotics over time observed in the same strain isolates shows that antibiotic tolerance of biofilm is a key pathoadaptation by persister isolates to the sinonasal environment of CRS patients.

Our study found that out of the 34 CI pairs, 14 (41%) were highly related strains based on a two-step approach considering their MLST/PopPUNK clustering and low core genome between isolate pair SNV. Past studies have posited that longitudinal follow-up of *S. aureus* nasal carriage in the healthy population colonisation by a single strain occurs between the 73% and 77% over time (Muthukrishnan et al., 2013). Additionally, Drilling et al. identified that 79% of recalcitrant CRS patients have a persistent *S. aureus* strain in their paranasal sinuses (Drilling et al., 2014). However, these results are based on MLST and pulsed-field gel electrophoresis typing, which may overestimate persister isolates due to less accuracy in discerning strains’ genetic relatedness compared to WGS. Thunberg et al. reported a lower proportion (20%) for single-strain long-term colonisation in CRS patients using WGS. This proportion is lower than the 41% identified in this study. A possible explanation for this might be a longer time between CI pairs collection (10 years) and a lower sample size (n=15) in that study (Thunberg et al., 2021). Furthermore, a considerable proportion of the isolates (35.2%) did not belong to any known CC based on MLST analysis, while 2 CIs pairs belonged to the same CC or VLKC while having relatively high SNV and SV counts between them, highlighting the limitations of this approach in characterising the genetic similarity of *S. aureus* populations.

The definition of closely related clonal isolates in the literature often employs a threshold-based approach using SNVs divergence. This is typically done by mapping short-read sequences to a reference sequence or calculating core genome SNPs. (Coll et al., 2020; Lagos et al., 2022). However, using long-read sequencing technologies enabled us to assemble near-perfect genome assemblies and plasmids instead, which facilitated using the first timepoint isolate as a reference for each longitudinal pair. This revealed that even in low SNV divergent isolate pairs, isolates undergo significant structural changes, such as prophage acquisition, mobile genetic element insertion or loss, and plasmid acquisition, that are difficult or impossible to capture using SNVs only.

Additionally, we found no relationship between the number of SNVs and the presence or number of structural variants. Combined with other sequential genomics studies that have revealed similar structural changes in the context of bacteraemia, we suggest that SNP-based cutoffs cannot fully capture CIs’ genomic adaptations and, instead, methods that take into account structural variation should complement the analysis (Giulieri et al., 2018; Giulieri et al., 2022).

MSCRAMM genes are known to be involved in epithelial adhesion and biofilm formation (Foster, 2019; Raafat, Otto, Reppschlager, Iqbal, & Holtfreter, 2019). While a single gene or gene cluster was not found to be indicative of colonisation, our comprehensive genomic analysis demonstrated that MSCRAMM genes frequently exhibited variability in persister strains over time, implying that selection pressure might act on the MSCRAMM genes in chronic colonisation. Interestingly, we found that persisters commonly had both small and structural MSCRAMM gene variants over time, suggesting that once colonisation has occurred, persister strains may attenuate their virulence profiles by adaptive evolution over time (Howden et al., 2023). Detailed analysis of the *sdr* locus deletion in 4875 revealed recombination of the folding domains from the *sdrC* gene and the wall-spanning and sort domain of the *sdrD* gene, suggesting that intra-host surface adhesin modulation can occur. To our knowledge, such recombination has not been previously reported in *S. aureus*. While it is known that serine-aspartate repeat MSCRAMM proteins are variable and contribute to biofilm formation (Ajayi et al., 2018; Barbu, Mackenzie, Foster, & Hook, 2014), more work needs to be done to characterise the relationship between divergent serine-aspartate repeat MSCRAMM proteins and their relationship to within-host adaptation.

We employed the Nanopore long-read Rapid Barcoding Kit library preparations, which have been demonstrated to retrieve small plasmids effectively (R. R. Wick, Judd, Wyres, & Holt, 2021). Our study revealed that *S. aureus* isolated from the sinonasal cavity of CRS patients frequently carried plasmids, regardless of their persistence. While our sample size was insufficient to establish a definitive association between plasmid carriage and lineage, we confirmed that these plasmids commonly contained the beta-lactamase gene *blaZ*, which has been frequently found in *S. aureus* strains since the advent of penicillin (Turner et al., 2019). Furthermore, our results suggest that *blaZ* encoding plasmids become more prevalent over time, but given the limited sample size, cautious interpretation is warranted.

Overall, our study revealed a trend of higher plasmid copy numbers in the long-read dataset compared to the short-read data, consistent with the findings of Wick et al. (R. R. Wick et al., 2021). We speculate that the discrepancy in copy numbers estimation in the long-read dataset compared to the short may be attributed to the PCR-free nature of the Rapid Barcoding Kit, which could potentially reduce bias compared to PCR-based short-read methods. However, this hypothesis requires further investigation, particularly as long-read sequencing becomes more commonly used, as there is scant literature on the impact of different library preparations on plasmid copy number estimation. Although limited knowledge exists regarding the fitness cost of carriage and copy number of plasmids for *S. aureus*, our study observed an increase in the copy number of conserved plasmids over time in the long-read dataset for the persistent isolates not in the short-read dataset.

Consistent with previous studies, we observed a high prevalence of macrolide resistance in our set of CIs isolated from CRS patients (Bhattacharyya & Kepnes, 2008). Additionally, our findings are consistent with previous studies that have shown a significant decrease in the effectiveness of antibiotics against *S. aureus* biofilms compared to their planktonic counterparts (C. W. Hall & Mah, 2017). Only doxycycline was found to have a strong ability to reduce biofilms to near eradication. However, this was only at concentrations exceeding the therapeutic window in humans. These results suggest that antibiotics alone may not be sufficient for eradicating *S. aureus* biofilms in the sinuses of long-term colonized CRS patients, as biofilms are a common feature in the sinuses of CRS patients (Foreman, Psaltis, Tan, & Wormald, 2010; Singhal, Psaltis, Foreman, & Wormald, 2010). Our finding of a frequent persistence of a single *S. aureus* strain in CRS patients is further evidence that the use of topical and systemic antibiotics alone may not be sufficient to eradicate the bacteria. However, we observed a substantial reduction of *S. aureus* biofilms for mupirocin in concentrations achievable when applied topically in saline-based irrigations (Kim & Kwon, 2016). Therefore, saline nasal irrigation mixed with mupirocin could play a role in the peri-operative phase of functional endoscopic sinus surgery of CRS patients by reducing the *S. aureus* biofilm, which has been correlated with delayed wound healing and poor post-surgical outcomes (Percival, 2017; Psaltis et al., 2008).

Pathoadaptation of persistent colonisers in (chronic) inflammatory conditions has been described for pathogens such as *S. aureus* and *Pseudomonas aeruginosa* (Howden et al., 2023; Rossi et al., 2021). A surprising finding in this study was the significantly increased biofilm antibiotic tolerance over time of the *S. aureus* strains that are persistent. This increased tolerance was correlated with an increase in the biomass of biofilms of the persister isolates. The biofilm production and viability in persister CIs were lower compared to the non-persister strains at the first timepoint. This suggests that CIs with attenuated biofilm production capacities are more likely to persist in the niche. It can be postulated that the observed increased antibiotic tolerance in those persistent strains over time assists them in their host adaptation, making them well-equipped to occupy and dominate the sinonasal microenvironment of CRS patients, which are frequently exposed to antibiotics. However, it is essential to note that the sinonasal cavity of humans is a relatively low-nutrient environment for bacteria, and high biofilm production might bring a fitness cost (Krismer et al., 2014). The increased biofilm production adaptation may arise during disease exacerbation with high bacterial load in the sinuses and antibiotic exposure. It may present a fitness cost during periods between exacerbations, allowing strains with less biofilm production to take over the niche. The data on non-persistent strains did not show a reduction in biofilm production between the first and second strains. However, the exact timepoint of strain change was not known.

Various mechanisms have been postulated to contribute to biofilm-based antibiotic tolerance of bacteria and the production of extracellular polymeric substances (C. W. Hall & Mah, 2017; Karygianni, Ren, Koo, & Thurnheer, 2020). Although we observed episode-specific mutations in the persistent isolates, we noted that genes involved in adhesion and biofilm formation were frequently affected, suggesting that the accumulation of mutations in different genes can lead to similar phenotypic adaptations.

### Limitations

The findings of this study have to be seen in the light of some limitations. Since the study was limited to *S. aureus* CIs isolated from patients suffering from CRS, it was not possible to compare the results to longitudinal CIs from carriers. Longitudinal CIs from carriers with extended follow-up are practically hard to obtain. Notwithstanding this limitation, this study offers some insight into the genomic and phenotypical adaption of *S. aureus* in the sinuses of CRS patients. Furthermore, the scope of the genomic analysis in this study was limited due to the low sample size. The genomic complexity of *S. aureus* does not lend itself to genome-wide association studies in low sample size populations. A natural progression of this work is to analyse the genome of specific CIs pairs and all the in-between CIs to identify a genomic target that might be involved in the phenotypical adaptation.

### Conclusion

Our findings provide insights into *S. aureus* persistence in difficult-to-treat CRS and highlight the resilience of bacterial biofilms. Our results shed light on the genomic and phenotypic changes associated with the persistence of *S. aureus* in chronically colonised CRS patients. Further studies are needed to understand the mechanisms underlying these adaptations and their potential survival benefit to identify potential targets for developing new eradication strategies.

## Abbreviations

AERD: aspirin-exacerbated respiratory disease
AMR: Antimicrobial Resistance
CARD: Comprehensive Antibiotic Resistance Database
CDS: coding sequences
CFU: Colony Forming Unit
CI: Clinical Isolate
CLSI: Clinical and Laboratory Standards Institute
CRS: Chronic Rhinosinusitis
CRSsNP: CRS without Nasal Polyposis
CRSwNP: CRS with Nasal Polyposis
CV: Crystal Violet
MFU: McFarland Units
MIC: Minimum Inhibitory Concentration
MIC50: MIC required to inhibit the growth of 50% of organisms
MIC90: MIC required to inhibit the growth of 90% of organisms
MLST: Multi-Locus Sequence Typing
NCTC: National Collection of Type Culture
NGS: Next Generation Sequencing
ONT: Oxford nanopore technology
PCR: Polymerase Chain Reaction
SNV: Single-Nucleotide Variant
SV: structural variants
VFDB: Virulence Factor Database
VLKCs: Variable-length k-mer clusters
WGS: Whole Genome Sequencing

## Tables

**Table S1.** Biosample accession number for each sample.

| **Biosample accession** | **Sample id** | **Host id and timepoint** |
| --- | --- | --- |
| **SAMN32360832** | C22 | 420_T0 |
| **SAMN32360833** | C265 | 1415_T0 |
| **SAMN32360834** | C13 | 1676_T0 |
| **SAMN32360835** | C80 | 1992_T0 |
| **SAMN32360836** | C295 | 2911_T1 |
| **SAMN32360837** | C52 | 3344_T0 |
| **SAMN32360838** | C9 | 3997_T0 |
| **SAMN32360839** | C3 | 4681_T0 |
| **SAMN32360840** | C67 | 4875_T0 |
| **SAMN32360841** | C72 | 5047_T0 |
| **SAMN32360842** | C45 | 5142_T0 |
| **SAMN32360843** | C224 | 5519_T0 |
| **SAMN32360844** | C222 | 5562_T0 |
| **SAMN32360845** | C241 | 5728_T0 |
| **SAMN32360846** | C100 | 276_T0 |
| **SAMN32360847** | C364 | 276_T1 |
| **SAMN32360848** | C320 | 420_T1 |
| **SAMN32360849** | C235 | 539_T0 |
| **SAMN32360850** | C318 | 539_T1 |
| **SAMN32360851** | C148 | 1170_T1 |
| **SAMN32360852** | C79 | 1170_T0 |
| **SAMN32360853** | C324 | 1415_T1 |
| **SAMN32360854** | C76 | 1676_T1 |
| **SAMN32360855** | C208 | 1992_T1 |
| **SAMN32360856** | C240 | 2911_T0 |
| **SAMN32360857** | C113 | 3344_T1 |
| **SAMN32360858** | C195 | 3357_T1 |
| **SAMN32360859** | C24 | 3357_T0 |
| **SAMN32360860** | C353 | 3997_T1 |
| **SAMN32360861** | C121 | 4009_T0 |
| **SAMN32360862** | C255 | 4009_T1 |
| **SAMN32360863** | C56 | 4681_T1 |
| **SAMN32360864** | C188 | 4747_T1 |
| **SAMN32360865** | C32 | 4747_T0 |
| **SAMN32360866** | C16 | 4784_T0 |
| **SAMN32360867** | C70 | 4784_T1 |
| **SAMN32360868** | C294 | 4875_T1 |
| **SAMN32360869** | C21 | 4986_T0 |
| **SAMN32360870** | C273 | 4986_T1 |
| **SAMN32360871** | C351 | 5047_T1 |
| **SAMN32360872** | C133 | 5060_T0 |
| **SAMN32360873** | C179 | 5060_T1 |
| **SAMN32360874** | C149 | 5142_T1 |
| **SAMN32360875** | C209 | 5308_T1 |
| **SAMN32360876** | C91 | 5308_T0 |
| **SAMN32360877** | C206 | 5328_T0 |
| **SAMN32360878** | C276 | 5328_T1 |
| **SAMN32360879** | C136 | 5390_T0 |
| **SAMN32360880** | C197 | 5390_T1 |
| **SAMN32360881** | C155 | 5448_T0 |
| **SAMN32360882** | C339 | 5448_T1 |
| **SAMN32360883** | C196 | 5469_T0 |
| **SAMN32360884** | C342 | 5469_T1 |
| **SAMN32360885** | C183 | 5485_T0 |
| **SAMN32360886** | C312 | 5485_T1 |
| **SAMN32360887** | C182 | 5503_T0 |
| **SAMN32360888** | C233 | 5503_T1 |
| **SAMN32360889** | C349 | 5519_T1 |
| **SAMN32360890** | C333 | 5562_T1 |
| **SAMN32360891** | C245 | 5631_T0 |
| **SAMN32360892** | C314 | 5631_T1 |
| **SAMN32360893** | C285 | 5647_T0 |
| **SAMN32360894** | C355 | 5647_T1 |
| **SAMN32360895** | C309 | 5728_T1 |
| **SAMN32360896** | C325 | 5767_T0 |
| **SAMN32360897** | C363 | 5767_T1 |
| **SAMN32360898** | C311 | 5911_T0 |
| **SAMN32360899** | C357 | 5911_T1 |

**Table S2.** Metadata characteristics of chronic rhinosinusitis subjects and corresponding clinical isolate collection.

| Characteristic | CRSwNP | CRSsNP | Total |
| --- | --- | --- | --- |
| Host subjects-N | 29 | 5 | 34 |
| Gender male-N | 12 | 2 | 14 |
| Age-year (± SD) | 51 (± 11) | 54 (± 21) | 52 (± 15) |
| Aspirin sensitivity-N | 6 | 0 | 6 |
| Asthmatic-N | 17 | 2 | 19 |
| Mean time between CI pair collection- days (± SD) | 565 (±398) | 558 (±362) | 563 (±387) |

CRSwNP, chronic rhinosinusitis with nasal polyps, CRSsNP, chronic rhinosinusitis without nasal polyps.

**Table S3.** Compilation of metadata characteristics of clinical isolate by relatedness classification and host characteristics.

| Characteristic | Same strain pairs | Different strain pairs | Total |
| --- | --- | --- | --- |
| Count-N (% of total) | 14 (41%) | 20 (59%) | 34 (100%) |
| Gender of host-Male-N | 5 | 9 | 14 |
| Age of host-year (± SD) | 57 (± 15) | 50 (± 16) | 52 (± 15) |
| Phenotype of host-CRSwNP- N | 14 | 15 | 29 |
| Aspirin sensitivity of host-N | 3 | 3 | 6 |
| Asthma status of host-N | 7 | 12 | 19 |
| Mean time between CI pair collection- days (± SD) | 629 (±474) | 518 (±320) | 563(±387) |

CRSwNP, chronic rhinosinusitis with nasal polyps.

**Table S4.** Frequency distribution of Staphylococcus aureus isolates’ Minimum Inhibitory Concentration (MIC)

| Antibiotic | Timepoint * | MIC (%) |  |  |  |  |  |  |  |  |  |  |  | MIC<sub>50</sub> | MIC<sub>90</sub> | Resistant (%) |
| --- | --- | --- | --- | --- | --- | --- | --- | --- | --- | --- | --- | --- | --- | --- | --- | --- |
|  |  | Concentration (mg/L) |  |  |  |  |  |  |  |  |  |  |  |  |  |  |
|  |  | <0.06 | 0.06 | 0.125 | 0.25 | 0.5 | 1 | 2 | 4 | 8 | 16 | 32 | >32 |  |  |  |
| Erythromycin | T0 | 0.00 | 0.00 | 0.00 | 0.00 | 0.00 | 14.71 | 47.06 | 17.65 | 5.88 | 5.88 | 2.94 | 5.88 | 2 | >32 | 21 |
|  | T1 | 0.00 | 0.00 | 0.00 | 0.00 | 0.00 | 2.94 | 38.24 | 26.47 | 14.71 | 2.94 | 5.88 | 8.82 | 4 | >32 | 32 |
| Doxycycline | T0 | 0.00 | 2.94 | 26.47 | 52.94 | 8.82 | 5.88 | 0.00 | 2.94 | 0.00 | 0.00 | 0.00 | 0.00 | 0.25 | 0.50 | 0 |
|  | T1 | 0.00 | 0.00 | 5.88 | 29.41 | 47.06 | 5.88 | 2.94 | 5.88 | 0.00 | 2.94 | 0.00 | 0.00 | 0.50 | 2 | 3 |
| Clindamycin | T0 | 0.00 | 0.00 | 2.94 | 52.94 | 35.29 | 5.88 | 0.00 | 0.00 | 0.00 | 0.00 | 0.00 | 2.94 | 0.25 | 0.50 | 3 |
|  | T1 | 0.00 | 0.00 | 0.00 | 23.53 | 58.82 | 5.88 | 0.00 | 0.00 | 0.00 | 0.00 | 0.00 | 11.76 | 0.50 | 2 | 12 |
| Augmentin | T0 | 0.00 | 0.00 | 0.00 | 5.88 | 14.71 | 38.24 | 32.35 | 2.94 | 0.00 | 5.88 | 0.00 | 0.00 | 4 | 8 | N. A |
|  | T1 | 0.00 | 0.00 | 0.00 | 2.94 | 8.82 | 23.53 | 29.41 | 11.76 | 5.88 | 5.88 | 8.82 | 2.94 | 4 | 16 | N. A |
| Gentamycin | T0 | 0.00 | 0.00 | 0.00 | 2.94 | 17.65 | 29.41 | 35.29 | 2.94 | 5.88 | 2.94 | 0.00 | 2.94 | 4 | 16 | 6 |
|  | T1 | 0.00 | 0.00 | 0.00 | 0.00 | 2.94 | 11.76 | 41.18 | 14.71 | 11.76 | 5.88 | 2.94 | 8.82 | 4 | 16 | 18 |
| Mupirocin | T0 | 85.29 | 2.94 | 8.82 | 0.00 | 0.00 | 0.00 | 0.00 | 0.00 | 0.00 | 2.94 | 0.00 | 0.00 | <0.06 | 0.125 | N. A |
|  | T1 | 67.65 | 0.00 | 29.41 | 2.94 | 0.00 | 0.00 | 0.00 | 0.00 | 0.00 | 0.00 | 0.00 | 0.00 | <0.06 | 0.125 | N. A |
| Clarithromycin | T0 | 0.00 | 0.00 | 0.00 | 0.00 | 0.00 | 8.82 | 58.82 | 14.71 | 0.00 | 5.88 | 2.94 | 8.82 | 2 | 32 | 18 |
|  | T1 | 2.94 | 0.00 | 0.00 | 0.00 | 0.00 | 14.71 | 32.35 | 23.53 | 5.88 | 2.94 | 2.94 | 14.71 | 2 | >32 | 26 |

\*T0 and T1 represent the first and second CIs of the longitudinal *S. aureus* pairs. The light-grey fill represents the intermediate
susceptible range, and the dark-grey fill represents the resistant range. MIC50 and MIC90 is the minimum inhibitory concentration
(mg/L) that inhibited 50 and 90% of the isolates, respectively. Antibiotics without shading have no interpretive MIC breakpoint accessible.

## Figure

**Figure S1.**
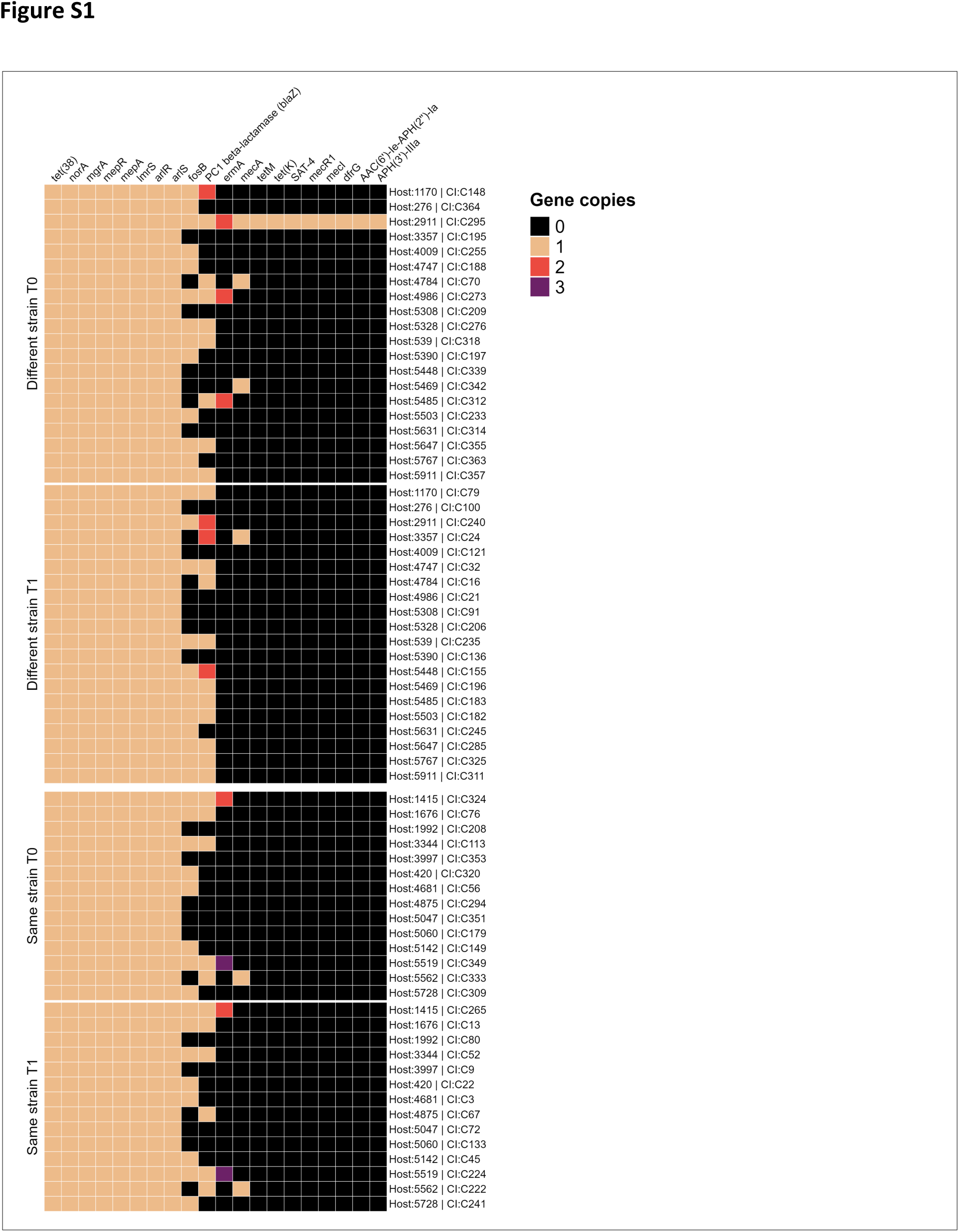
Gene copy matrix of antimicrobial resistance genes in the genomes of all *S. aureus* clinical isolates (n=68). The isolates are represented by the rows and grouped by strain-relatedness classification and the order of CI collection. The gene copies of antimicrobial resistance genes are indicated by colour.

**Figure S2.**
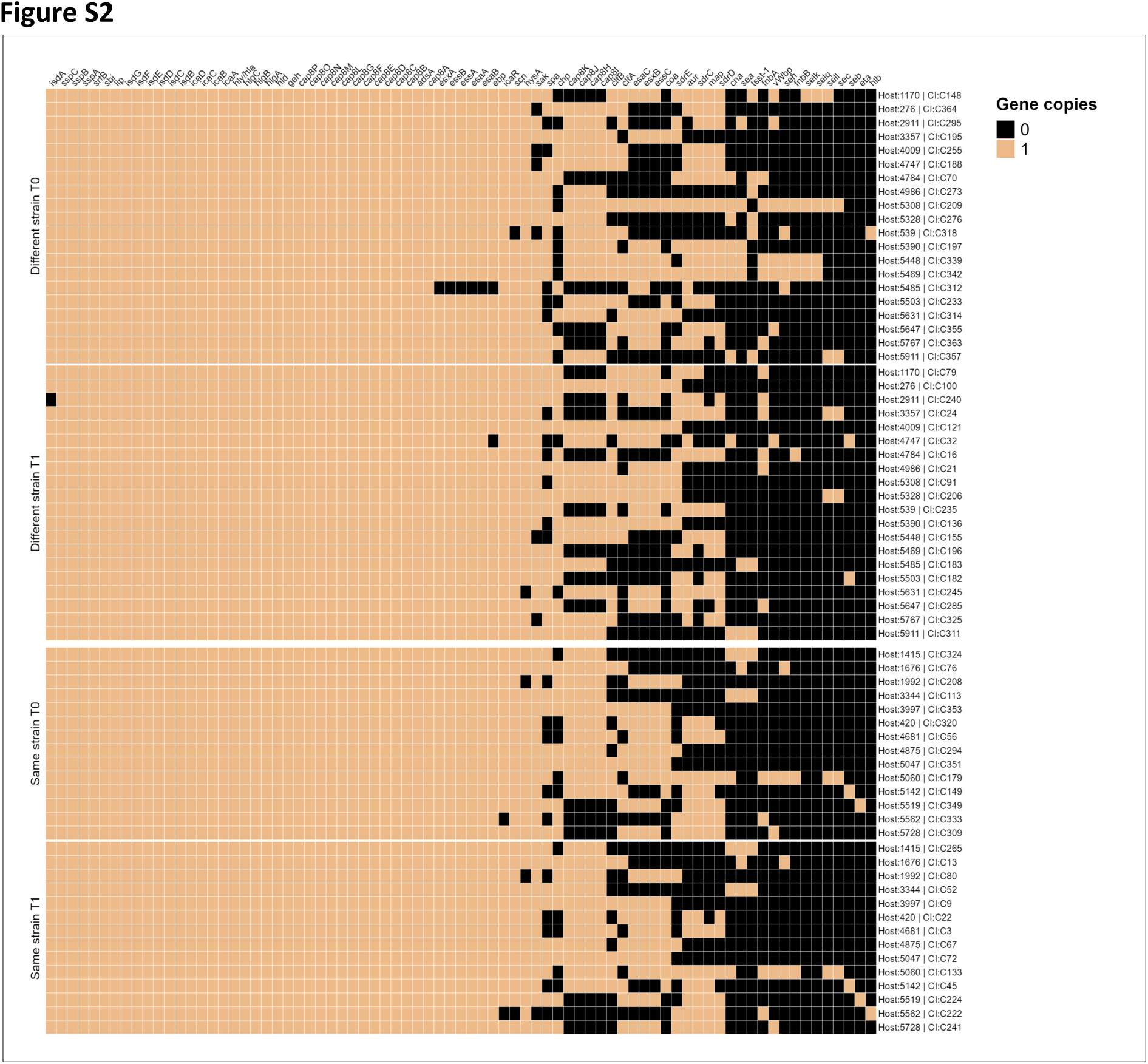
Matrix displaying the virulence genes in the genome of all CIs (n=68). CIs are grouped by strain relatedness and the collection sequence. The matrix is split into strains classified as ’same strain’ pairs (N=14) and ’different strain’ pairs (N=20). T0 and T1 indicate the first and second CIs groups of the sequential *S. aureus* pairs, respectively.

**Figure S3.**
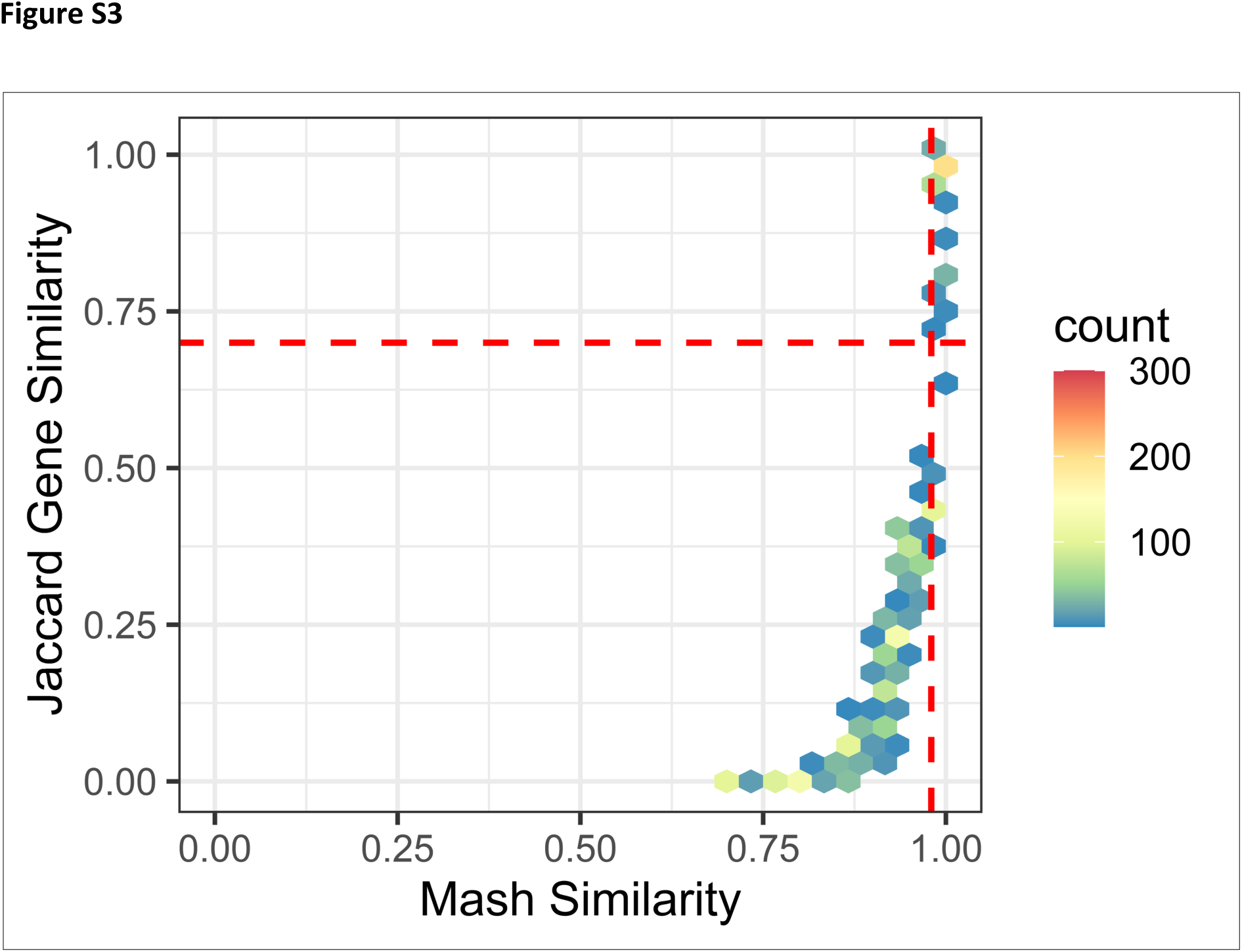
Hexagonal binned plot displaying the relationship between all the plasmids detected based on gene presence and absence and M similarity (n=2,756). The hexagonal cell colour represents the number of data points observed in that cell. Plasmids were considered the same using a threshold of Mash similarity distance greater than 0.98 and Jaccard index of gene presence and absence greater than 0.7.

**Figure S4.**
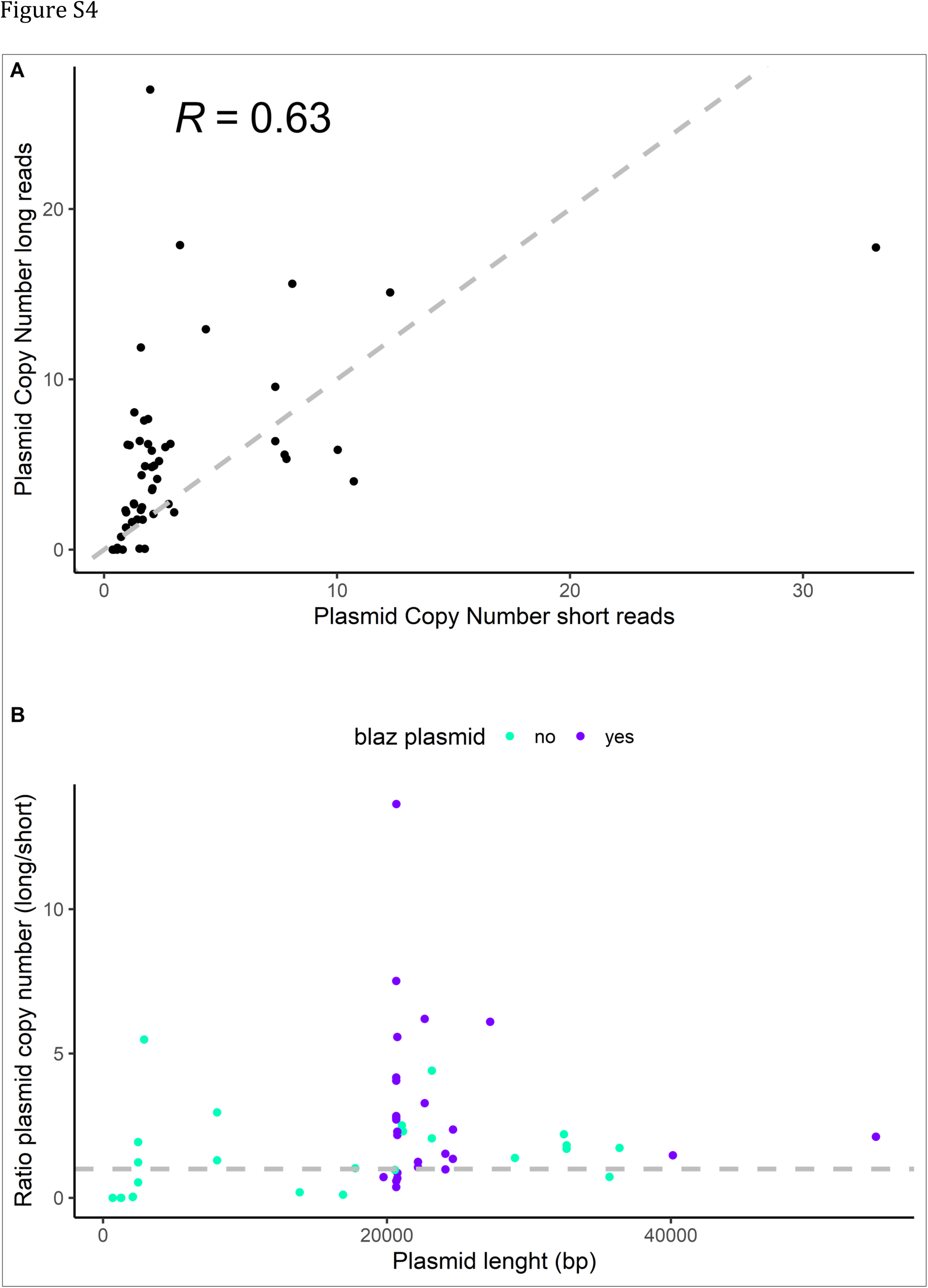
Comparison of plasmid copy numbers estimated from long-read sequencing versus short-read sequencing. (A) Each point represents a single plasmid (n=53), with the x-axis indicating the copy number from short-read sequencing and the y-axis indicating the copy number from long-read sequencing. The Spearman correlation coefficient is shown in the top left. (B) The ratio of the plasmid copy number (short/long) for all plasmids detected (N=53). *Blaz*-positive plasmids are indicated by colour. The grey line is the intercept of the ratio=1.

**Figure S5.**
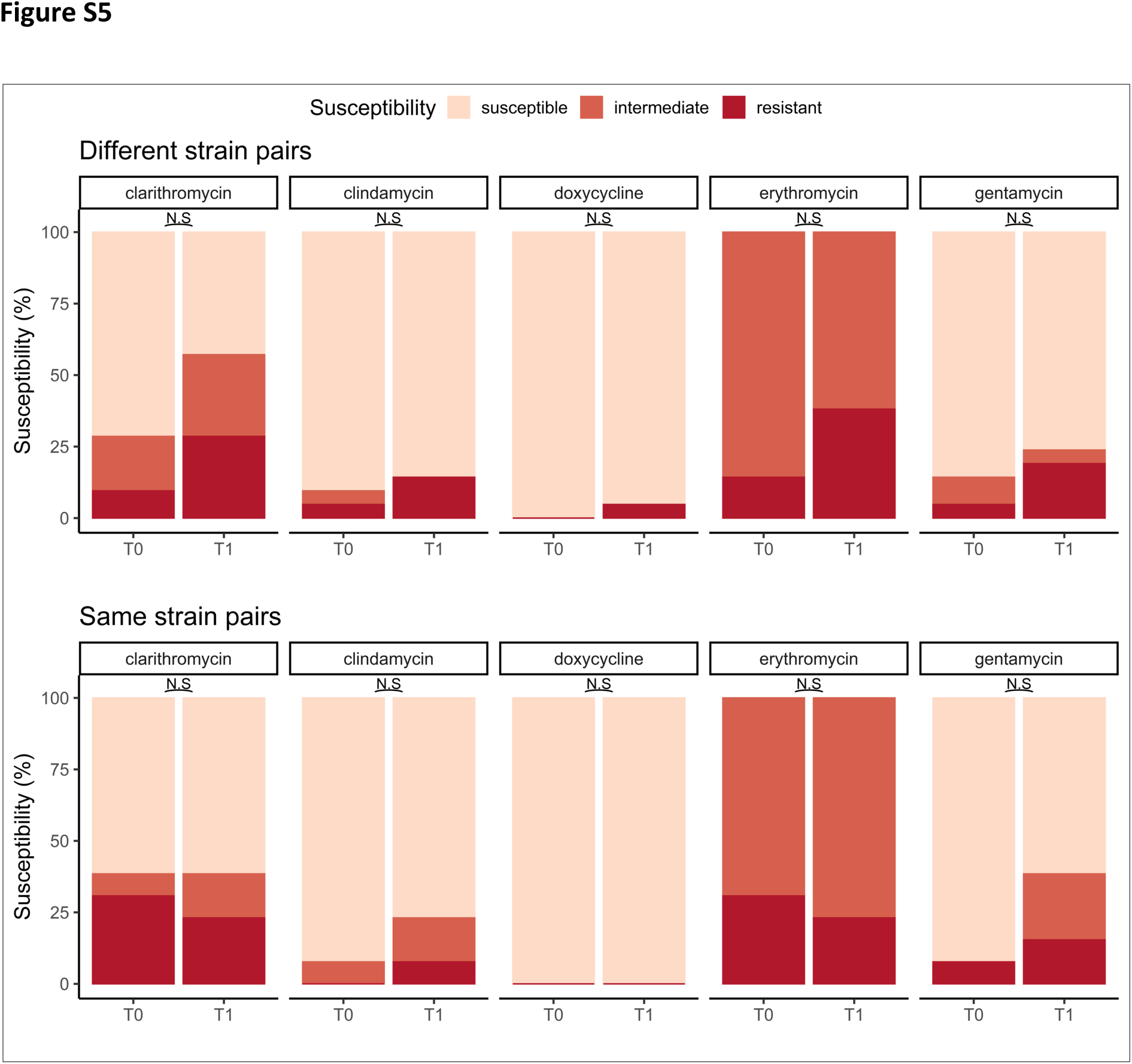
Planktonic antibiotic susceptibility of *S. aureus* isolates (n=68). Antibiotic susceptibility of *S. aureus* isolates is presented based on minimum inhibitory concentration (MIC) breakpoints adapted from the CLSI for isolates classified as the same and different strains (N=14, N=20). Fisher’s exact test was used to determine the significant difference in the proportion of resistant and non-resistant isolates between the T0 and T1 groups, with a threshold of p<0.05. MIC breakpoints are not available for augmentin and mupirocin.

**Figure S6.**
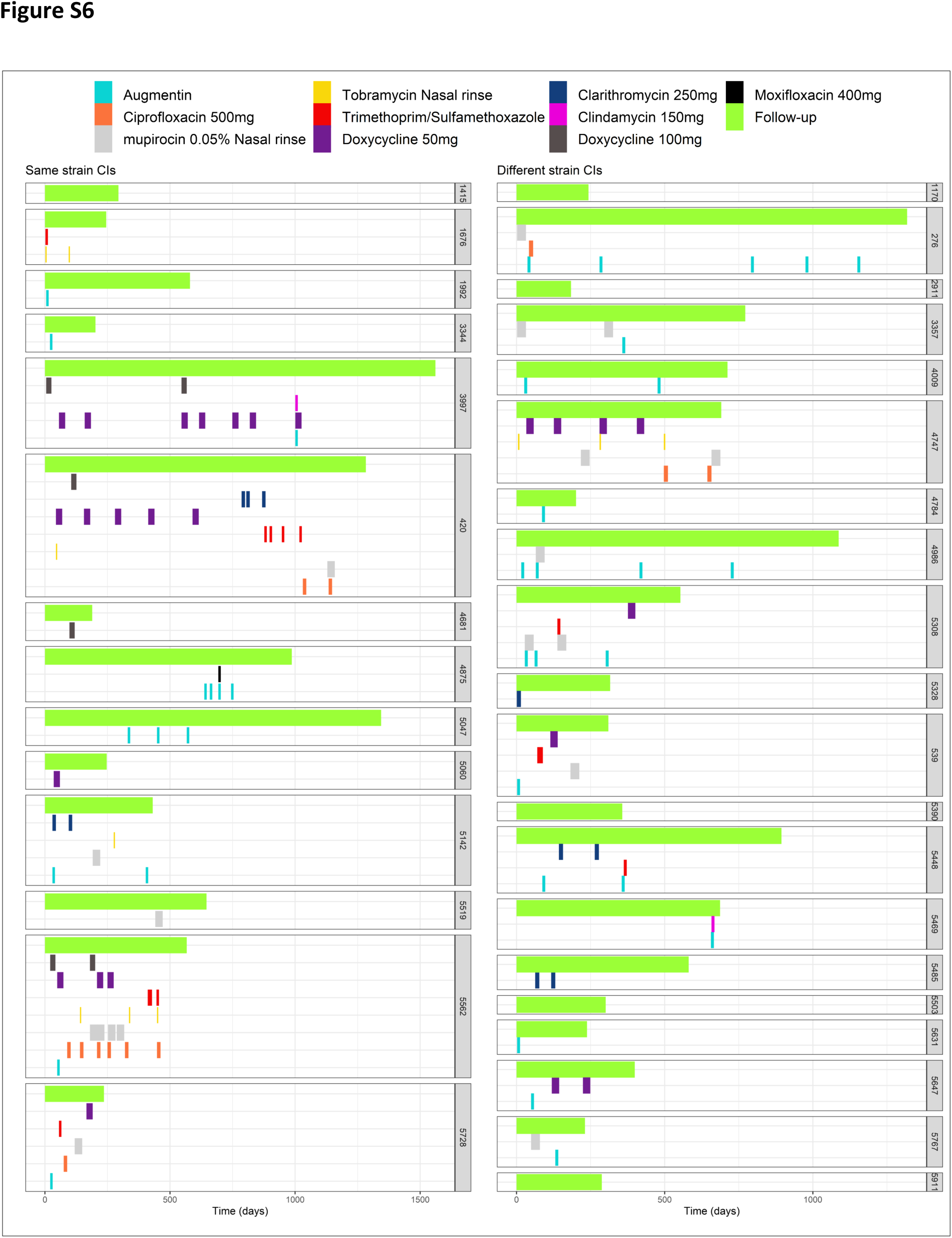
Swimmer plot showing the timeline of antibiotic prescription for each subject between the collection of the first and second timepoint isolates. The antibiotics are indicated by colour. The top green bar represents the time between isolate collection. The left column represents all ’same strain’ isolates, and the right column represents ’different strain’ isolates. Host ID numbers are indicated on the right side of the columns.

**Figure S7.**
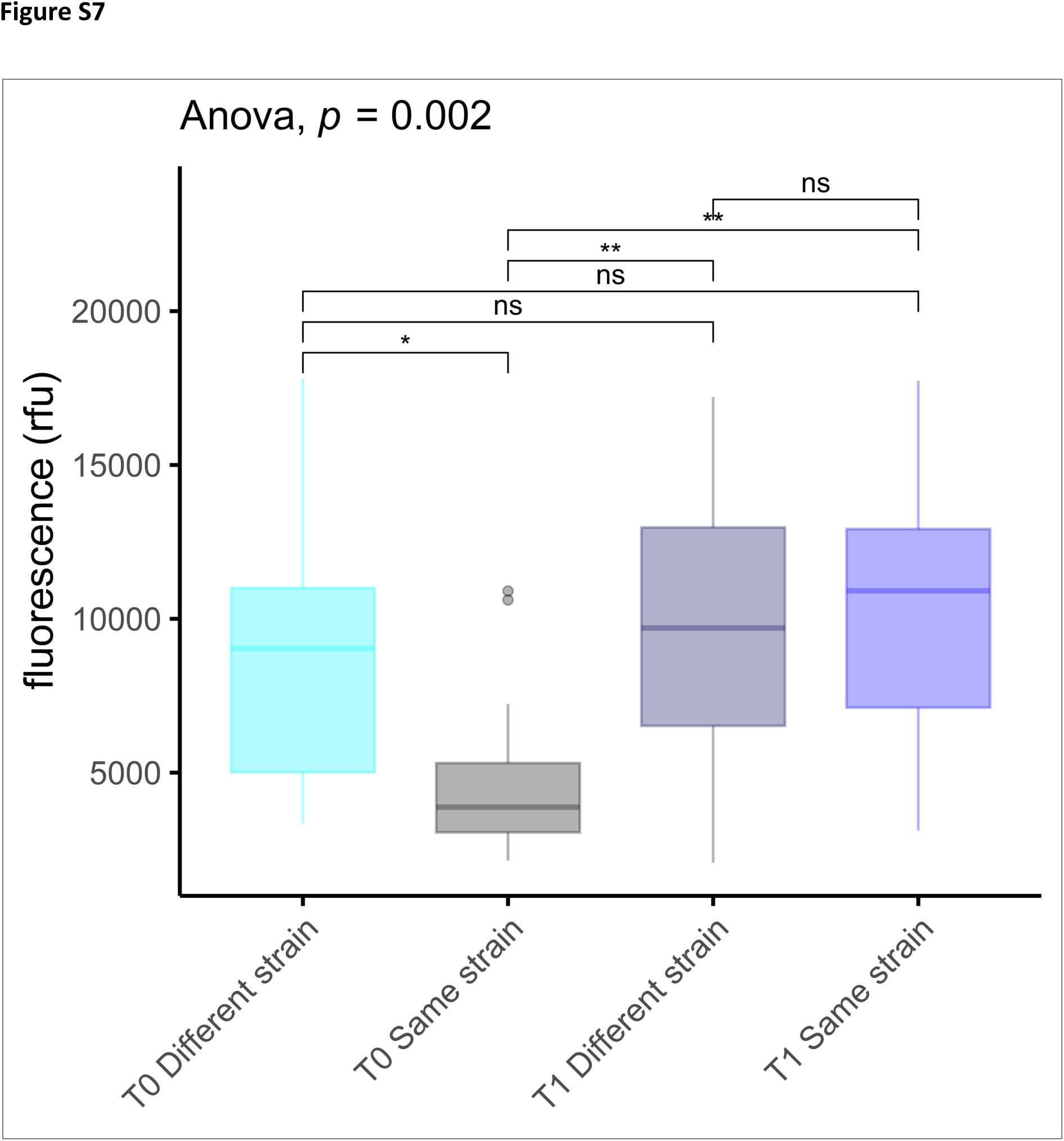
*S. aureus* biofilms viability after 48 hours of growth for all 68 clinical isolates (CIs). The x-axis indicates the first and second CIs classified as the ’same strain’ and ’different strain’. Significance was tested using ANOVA, and post-hoc pairwise t-test with Bonferroni correction applied for multiple comparisons.

## Supplementary text

### ST1

Chronic rhinosinusitis (CRS) diagnosis criteria as described by the EPOS: The presence of two or more symptoms, one of which should be either nasal blockage or nasal discharge with facial pain/pressure or loss of smell. The symptoms should last for more than 12 weeks. Patients were considered difficult-to-treat if no acceptable level of control was achieved despite adequate surgery, intranasal corticosteroid treatment and short courses of antibiotics or systemic corticosteroids in the preceding year of collection.

Asthma status and aspirin sensitivity were collected via self-reported questionnaires at the time of consent for the biobank. Furthermore, an ENT surgeon added the CRS subtype to the biobank after endoscopic assessment.

## Notes

**Conflicts of interest** The authors state that the study was conducted without any commercial and financial relationship that could be interpreted as a potential conflict of interest.

https://github.com/gbouras13/CRS_Saureus_Evolutionary_Landscape

### Competing Interest Statement

The authors have declared no competing interest.

